# Profiling diverse sequence tandem repeats in colorectal cancer reveals co-occurrence of microsatellite and chromosomal instability involving Chromosome 8

**DOI:** 10.1101/2020.12.23.422767

**Authors:** GiWon Shin, Stephanie U. Greer, Erik Hopmans, Susan M. Grimes, HoJoon Lee, Lan Zhao, Laura Miotke, Carlos Suarez, Alison F. Almeda, Sigurdis Haraldsdottir, Hanlee P. Ji

## Abstract

Colorectal carcinomas **(CRCs)** which have lost DNA mismatch repair display hypermutability evident in a molecular phenotype called microsatellite instability **(MSI)**. These mismatch repair deficient tumors are thought to lack widespread genomic instability features, such as copy number changes and rearrangements. To identify MSI for clinical diagnosis, current molecular testing looks for changes in mononucleotide or dinucleotide repeats. However, microsatellites have other types of sequence tandem repeats such as tri- and tetranucleotide motifs. These additional classes of microsatellites are generally not examined for MSI but are known to be unstable in a phenotype known as elevated microsatellite alterations at selected tetranucleotide repeats, or **EMAST**. We developed a sequencing approach that provides ultra-high coverage (>2500X) of microsatellite targets and cancer genes for profiling genomic instability. We assessed the diverse repeat motifs across 200 microsatellites. Our approach provides highly sensitive detection of MSI with high specificity, evaluates copy number alterations with high accuracy, delineates chromosomal instability **(CIN)** classification and deconvolutes subclonal architecture. By examining both MSI and CIN, we discovered mutations and copy number alterations that defined mixed genomic instability states of CIN and MSI, which are normally considered exclusive. An increase in copy number of chromosome arm 8q was prevalent among MSI tumors. Moreover, we identified an inter-chromosomal translocation event from a CRC with co-occurrence of MSI. Subclonal analysis demonstrated that mutations which are typically considered to be exclusive in MSI, shows mutual occurrence in MSI tumors with more sensitive characterization. Our approach revealed that *MSH3* mutations are a potential source of mixed genomic instability features. Overall, our study demonstrates that some colorectal cancers have features of both microsatellite and chromosomal instability. This result may have implications for immunotherapy treatment.

## INTRODUCTION

Microsatellites are composed of short tandem repeats **(STRs)** and are present throughout the human genome. STRs have different classes of motifs that include mono-, di-, tri- and tetranucleotide sequences. In colorectal carcinoma **(CRC)**, somatic mutations or methylation of DNA mismatch repair **(MMR)** genes (i.e. *MSH2*, *MLH1, PMS2, MSH6*) lead to increased mutation rates, particularly in microsatellites. Lynch syndrome is an autosomal dominant genetic disorder in which affected individuals are carriers of deleterious germline mutations in the MMR genes and have a substantially increased risk of CRC, as well as other malignancies. Somatic inactivation of the remaining wildtype allele of an MMR gene leads to inactivation of this DNA repair pathway and increased risk of developing tumors. Tumors with MMR loss display hypermutability in microsatellite sequences. This phenomenon is referred to as microsatellite instability **(MSI)** and is characterized by the accrual of insertions or deletions **(indels)** in either coding or non-coding microsatellite sequences. Based on specific criteria, tumors with high levels of microsatellite mutations are referred to as MSI-high **(MSI-H),** with mutation rates that are orders of magnitude greater than what is observed in tumors that are microsatellite stable **(MSS)** [1]. Importantly, MSI occurs in all classes of microsatellite repeats. However, nearly all published studies have exclusively focused on the presence of microsatellite mutations within mono- and dinucleotide repeats to assess MSI. Generally, there has not been a careful examination of other microsatellite classes, potentially missing important features of MSI and the underlying genomic complexity of these tumors.

There are a number of methods used for detecting MSI in cancer. One approach involves immunohistochemistry **(IHC)** staining of tumor sections for MMR protein expression of MLH1, MSH2, MSH6 and PMS2. A tumor lacking expression of one of these proteins is considered to have MSI. The most common molecular genetic assay for identifying MSI-H tumors requires PCR amplification of a limited panel of microsatellite markers. The MSI PCR test uses a multiplexed amplicon assay which requires testing five or more microsatellite markers – typically these are either mono- or dinucleotide repeats [2, 3]. Using capillary electrophoresis **(CE)**, tumor-specific changes in the microsatellite amplicon size indicate MSI when compared to the microsatellite genotypes from matched non-tumor cells. If a sufficient number of microsatellites demonstrate an allelic shift in size (e.g. two or more), the tumor is classified as MSI-H. PCR testing is considered to be the gold standard test for MSI-H. In comparison to MSI PCR, MMR IHC misses approximately 10% of tumors with MSI [4, 5]. Despite its important diagnostic role, MSI PCR tests have issues. PCR amplification leads to artifacts related to additional indels in microsatellites. This artifact is referred to as a stutter and complicates the identification of MSI, particularly when i) the change in a microsatellite allele occurs in a smaller fraction of the cells, and ii) the allelic shift in size is less than 3bp.

Next generation sequencing approaches for detecting MSI are based on targeted assays that enrich or amplify exon sequences (e.g. exomes and gene panels) or microsatellites [6–10]. When using targeted sequencing with gene panels and exomes, the presence of indels within exon-based mono- or dinucleotide repeats determines MSI status. Sequencing demonstrates the presence of mutations within microsatellite tracts, leading to allelic shifts and somatic genotypes. Generally, MSI NGS assays have high concordance with MSI PCR tests [6, 8–10]. However, MSI NGS tests are also susceptible to artifacts related to sequencing library amplification. PCR amplification stutter is also seen in all types of sequencing data. Therefore, detection of MSI at low tumor cellular fraction remains a challenge for NGS detection.

The traditional criteria for defining MSI is restricted to mono- or dinucleotide repeats. However, there is another category that involves elevated microsatellite alterations at selected tetranucleotide repeats **(EMAST)**. This category of microsatellite alterations is thought to be related to changes in the function of *MSH3*, another gene of the MMR pathway. MSH3 loss of function is characterized by instability in dinucleotide or longer repeats [11]. CRCs with EMAST have been reported in up to 50% of tumors; EMAST positive CRCs have changes in a variety of pathways that lead to genome instability (e.g. MSI, CpG island methylator phenotype, etc.) [11]. The EMAST phenotype may be associated with an elevated microsatellite mutation rate of mono- and dinucleotide repeats, but this is not consistently observed [12–15]. EMAST CRCs are also frequently associated with chromosomal instability **(CIN)** where portions of the genome show copy number alterations, aneuploidy and rearrangements [11]. However, there are few, if any, genomic studies that have examined these EMAST CRCs in detail. Moreover, there are no reports of EMAST tumors analyzed with whole genome sequencing **(WGS)**.

Detection of EMAST involves testing a set of tetranucleotide microsatellites for instability changes. The most common method involves PCR genotyping assays analyzed with CE [12]. Currently, there are no established markers or criteria which are used in classifying EMAST [16]. Recent studies have relied on using five or more tetranucleotide markers; a tumor is considered to be EMAST positive when 30% or more of the markers are unstable compared to the matched normal DNA genotypes [12–15]. NGS methods for detecting EMAST are generally not available, seeing that most targeting assays do not include microsatellites, such as tetranucleotide repeats [6–9]. Generally, exons lack tetranucleotide repeats of sufficient length and as a result, exome or targeted gene sequencing will miss these genomic instability features. Further, targeted sequencing assays with short reads may not span the entire length of tetranucleotide microsatellites, which are frequently longer than mono- and dinucleotide repeats.

Determining the MSI status of CRCs and other tumors is of increasing importance given advances in cancer immunotherapy. MSI-positive CRCs respond to immune checkpoint therapy while CRCs with CIN do not. MSI-related indels in exons produce frameshift mutations within a gene, leading to a higher number of novel peptides, also referred to as neopeptides, from the translated mutated protein. For MSI-H cancers, these neopeptides provide an abundance of unique cancer antigens that are absent from normal colon and rectal cells. Pembrolizumab and other immune checkpoint drugs stimulate the immune system such that T cells recognize these highly immunogenic cancer cells and their cancer-related neoantigens [17]. Thus, the patient mounts an immune response against his or her own cancer. Given the therapeutic predictive nature of MSI status for immunotherapy, molecular genetic testing is a diagnostic requirement for receiving immune checkpoint therapies. Surprisingly, anywhere from 10 to 40% of MSI-H tumors do not show a response to immunotherapy. Even when there is a response as seen in progression free survival, up to one third of patients with MSI-H tumors show resistance to immunotherapy, continued tumor growth and mortality related to their metastasis [17–19]. Therefore, it remains an open question as to why a significant proportion of MSI-H tumors do not respond or progress while on immune checkpoint therapy.

As noted, nearly all studies determining the presence of MSI in CRCs examine only mono- and dinucleotides. The same holds true for other cancer types with MSI [20]. Limiting the evaluation of MSI to only two classes of microsatellites overlooks more complex genetic features, such as instability in tetranucleotide repeats. Addressing this gap in our knowledge about cancer mutations in microsatellites, we developed a new sequencing approach to profile instability across different classes of microsatellites and cancer genes. Our analysis included an expanded set of mono-, di-, tri- and tetranucleotide repeats with minimal amplification error and high read coverage. We used ultra-high depth sequencing coverage in the thousands for sensitive and specific detection of somatic events such as microsatellite and gene mutations. Thus, one can quantify genomic changes that are indicative of genetic heterogeneity and subclonal diversity present in a given tumor. Simultaneously, we detected CIN via the high-accuracy identification of copy number alterations in cancer genes.

We profiled diverse microsatellite classes while considering additional features of genomic instability that may affect cancer driver genes. As we demonstrate on a cohort of CRCs, this approach provided quantitative microsatellite profiles informative for MSI and EMAST, revealed mixed classes of CIN and delineated genetic heterogeneity indicative of subclonal structure. Our profiling results showed that some colorectal cancers have both MSI and CIN co-occurrence. To further characterize the extent of chromosomal changes, we conducted WGS on tumors with mixed genome instability features. Whole genome analysis confirmed our findings from the targeted approach and showed evidence of an inter-chromosomal rearrangement in MSI CRCs. Changes in Chromosome 8 structure occurred frequently and were validated in other genomic data sets. Our results may have potential clinical implications for the evaluation of MSI CRCs for immunotherapy.

## METHODS

### Cancer samples

This study was conducted in compliance with the Helsinki Declaration. All patients were enrolled according to a study protocol approved by the Stanford University School of Medicine Institutional Review Board (IRB-11886 and IRB-48492). Informed consent was obtained from all patients. Tissues were obtained from the Stanford Cancer Institute Tissue bank and the Landspitali University Hospital.

Matched tumor and normal specimens underwent histopathology review to mark areas of tumor and normal tissue on hematoxylin and eosin-stained tissue sections and on the corresponding paraffin blocks. The samples were generally 60% tumor purity or higher. We macro-dissected the samples to provide improved tumor purity and extracted genomic DNA from the matched normal and tumor CRC samples. The dissected tissue was homogenized and processed using the E.Z.N.A. SQ RNA/DNA/Protein Extraction Kit (Omega Biotek Inc.). Briefly, we lysed the cells using the provided lysis buffer (SQ1); precipitated and removed proteins with protease (SQ2) and NaOAc; precipitated nucleic acids with isopropanol; washed; and re-suspended nucleic acid pellets in 10 mM Tris-HCl (pH 8.0) buffer. We removed RNA species in the nucleic acid via the addition of 4 μg of RNase A (Promega), and incubation at 37°C for 1 hour. After incubation, each sample was purified with AMPure XP beads in a bead solution-to-sample ratio of 1.5. Nucleic acids were quantified pre- and post-RNase treatment using a Thermo Scientific NanoDrop™ 8000 spectrophotometer or Qubit Broad Range DNA kit (Thermo Fisher Scientific, Waltham, MA). In addition, some genomic DNA was obtained from punched tissue cores using Maxwell FFPE tissue LEV DNA purification kit (Promega) per the manufacturer’s guidelines and quantitated based on the same protocol as described.

### Joint sequencing of microsatellites and cancer genes

We used a targeted sequencing technology which provides ultra-deep coverage and enables amplification-free libraries with reduced PCR error **(Figure 1a)** [21–23]. Referred to as oligonucleotide-selective sequencing **(OS-Seq)** this assay uses only a single primer, also called primer probe, that anneals to a target sequence. As a result, this method avoids issues found with traditional PCR or bait-hybridization enrichment of target exons **(Additional File 1: Figure S1)**. Extension from the target-specific primer copies the target sequence in a massively multiplexed fashion. Primer design, library preparation and sequencing are described in full details per the **Supplementary Methods (Additional file 1)**. We designed primer probes for 225 microsatellite markers and 1,387 exons from 85 genes **(Additional File 2: Table S1; Additional File 3)**. The microsatellites included diverse STR motifs **(Additional File 2: Table S2)**. These primers target mono- and dinucleotide repeats, as well as tri- and tetranucleotide repeats; the broader representation of motifs allowed us to distinguish different types of MSI, including those markers used for MSI PCR testing and EMAST. Using results from the Cancer Genome Atlas project **(TCGA)**, we selected the cancer genes with the highest frequency of mutations present in colorectal and gastric cancer [1, 24]. The targeted exons from 85 genes were located across all autosomes and the X chromosome **(Additional File 2: Table S3)**.

**Figure 1.**
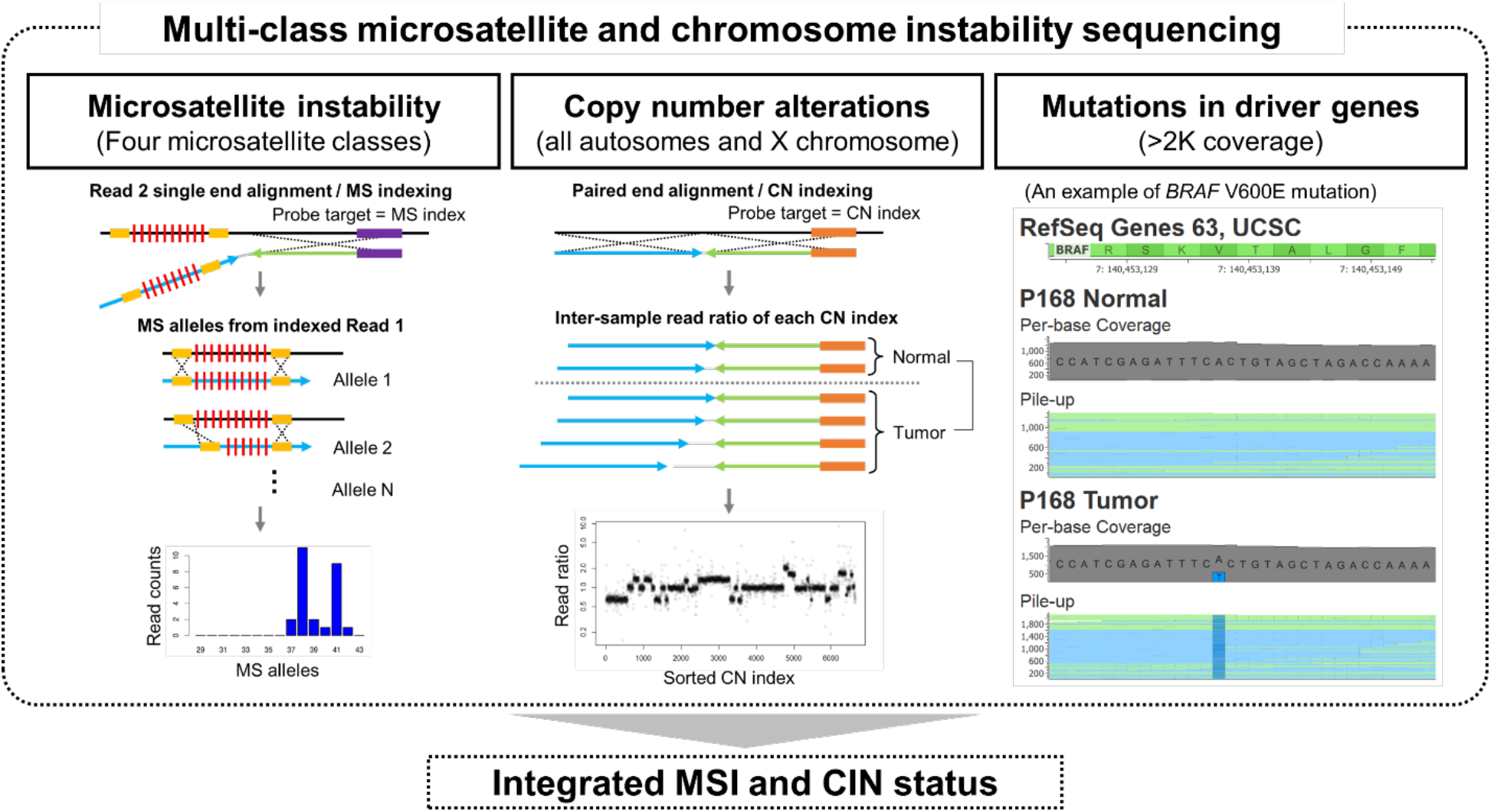
An integrated sequencing-based determination of microsatellite and chromosomal instabilities. An overview of the sequencing analysis that simultaneously determines microsatellite instability **(MSI)** and chromosomal instability **(CIN)** are shown. Mutations in driver genes also provide supplementary information that further supports the integrated determination. To determine MSI status based on four classes of microsatellites (mono-, di-, tri- and tetranucleotide repeats), after single end alignment of Read 2 sequences, the reads are indexed if aligning on target. The indexing informs the repeat motif and flanking non-repeat sequences, which are expected to be sequenced in the corresponding Read 1 sequence. We count reads for every observed microsatellite allele (i.e. number of motif repetitions), and generate a histogram, of which the comparison between tumor versus normal samples determines somatic microsatellite mutation. To determine CIN status, we analyze copy number alterations. After paired end alignment, the reads are indexed if aligning on target. An inter-sample comparison of read counts sharing the same index determines copy number alteration if the ratio is unusually high or low compared to the average ratio from all the indexes. Additionally, the targets for copy number analysis are also well-known driver genes. Our ultra-deep PCR-free targeted sequencing enables sensitive and quantitative detection of pathogenic mutations, and the mutation profile matching the molecular subtype improves the validity of our methods. Shown here is an example of a BRAF V600E mutation detected from an MSI tumor.

### Microsatellite mutation calling

We developed a bioinformatics pipeline to identify somatic changes to microsatellites [23]. Our analysis leveraged the unique properties of having the targeting primer sequenced as well as the genome target. The script and required data files are available at ‘https://github.com/sgtc-stanford/STRSeq’. An overview of the microsatellite genotyping process is illustrated in **Figure 1a**. Briefly, after single-end alignment to the NCBI v37 human genome, we added a custom SAM file tag **(ZP)** to the aligned reads. This tag denotes which microsatellite probe generated the read as well as the linked target STR information (e.g. repeat motif and flanking sequences). We used the Read 2 **(R2)** aligned sequences, which include the capture probe sequence and residual genomic sequences, to determine the tag for paired end reads. This indexing method does not require Read 1 **(R1)** sequences to align to the genome; both aligned and unaligned reads are tagged based on alignment of their R2 mate to a designated primer probe sequence. After the indexing, we evaluate reads to determine whether both the expected 5’ and 3’ STR flanking sequences are present in R1. For this study, we aggregated the ‘STR-spanning’ reads, which contain a combined sequence that includes an 8-base 5’ flanking sequence, a variable length region containing at least a minimum number of STR motif repeats, and an 8-base 3’ flanking region. Using the reads that contain this combined sequence, the STR motif repeat count was calculated by dividing the number of bases in the variable length region by the length of the STR motif. For example, if the variable length region is 28 bases and the STR motif is GATA (tetranucleotide), then the STR motif repeat count is seven. The R1 reads encompassing entire STRs (i.e. STR-spanning reads) are counted and summarized by motif repeat count (i.e. allele) to provide a basis for determining the genotype and extent of mutations in microsatellites. The distribution of alleles and relative percentage of reads for each allele was used to compare tumor versus normal samples **(Additional File 1: Figure S2a)**.

For a given microsatellite locus, we let *y* = (*y*_1_, …, *y*_*n*_) be the read count for each of the *n* alleles. Letting *S* = ∑_*i*_ *y*_*i*_ be the total number of reads observed for the locus, we computed the allele coverage proportion vector for the microsatellite as follows:

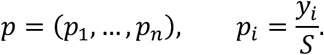

These proportion vectors exhibited peaks at alleles that are present in a sample. Using samples from a matched normal source, we first determined alleles *A_N_* based on the criteria previously described [23]. We assumed n(*A_N_*) ≤ 2. To determine whether a microsatellite locus has a somatic allele shift in tumor DNA *T* versus its paired normal *N*, we used a weighted Euclidean distance between the two proportion vectors:

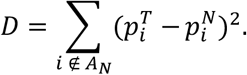

We excluded the proportions of normal alleles, *A_N_*, when calculating *D* (i.e. a zero weight for normal alleles). This weighting minimized the contribution of copy number changes in the distance. If *D* is greater than a given threshold, we consider that the microsatellite locus has an allele shift. The threshold was decided to minimize false positive and negative rates after fitting the most probable Gaussian mixture densities to our data **(Additional File 1: Figure S2b)**. To evaluate the extent of MSI of a tumor, we calculated the fraction of microsatellite loci with allele shift versus the total number of measured microsatellite loci.

### Microsatellite PCR genotyping

For mononucleotide repeat instability, we used the PowerPlex MSI analysis system v1.2 (Promega) to test five markers following the manufacturer’s protocol. For EMAST, we used a set of primers that amplify tetranucleotide microsatellites (D20S82, D20S85, D8S321, D9S242, and MYCL1) as has been previously published **(Additional File 2: Table S4)** [12]. The 12.5-μl reaction included two ng of gDNA, 0.4 μM of each primer, 1X Buffer I with MgCl_2_, 0.2 mM of each dNTP, and 1.25 units of AmpliTaq Gold polymerase (Thermo Fisher Scientific). The samples were denatured at 95°C for 5 min, followed by 35 cycles of 30 sec at 95°C, 90 sec at 58°C, and 30 sec at 72°C. The final steps for amplification involved an incubation at 72°C for 10 min and cooling to 4°C. We used ABI 3130xl Genetic Analyzer (Thermo Fisher Scientific) using a recommended matrix standard setting. PowerPlex 4C (Promega) and DS-33 (Thermo Fisher Scientific) matrix standard kits were used for the mono- and tetranucleotide repeat assays, respectively. Using the raw signal data (fsa files), the Peak Scanner v1.0 (Thermo Fisher Scientific) program provided the size of detected amplicons. We used a criterion where a shift in allele size of 3bp or more in the tumor sample compared to matching normal sample indicated instability at a given microsatellite locus.

### Identifying copy number alterations

We used the data from paired-end alignment to perform copy number analysis and somatic mutation calling. As with MSI analysis, the ‘ZP’ sam tag is added to both R1 and R2 reads. The number of reads sharing each primer ID is determined by counting ‘ZP’ tags, and represents how many DNA molecules are captured by the primer. Therefore, by comparing the per-probe read counts between tumor and normal samples, copy number changes can be measured. The per-probe read counts were first normalized by the total number of reads from all the probes, and then the log2 ratio between tumor versus normal was calculated. When calculating the ratio, we excluded the probes that showed high variation from normal DNA as determined by a Z-score. Specifically, the probes having a Z-score greater than 2 or less than −2 were excluded. In addition, we corrected systemic biases of the log2 ratios in terms of GC% of probe capture sequence and the per-probe read counts. The adjustments were made by the locally weighted scatterplot smoothing method. For example, the values at the regression line were set to zero. Using the per-probe ratio, which were normalized and corrected, median ratios for all the target genes were calculated as a representation of copy number. These values were used for generating heat maps as well. The per-gene median ratios were also used in comparison with WGS ratios.

### Copy number classification

We used an extreme gradient boosting algorithm called XGBoost to train a multi-class model and make predictions about the copy number classification [25]. The XGBoost method was implemented in the R package xgboost. We used the copy number data from the CRC TCGA study as the training set. For all analysis, we used the softprob objective function and ran it 100 times. Other training parameters such as eta, gamma, minimum child weight and maximum depth were all set as default values. An XGBoost model was then trained on the copy number cluster labels, using the 82 signature genes as features from the targeted sequencing assay.

### Identifying gene mutations

Paired-end read alignment was used for detection of somatic mutation. We used the Sentieon TNseq package (v201808.03; Sentieon Inc, Mountain View, CA) to preprocess the alignments and to call somatic variants following the best practice guidelines. Sentieon TNseq consists of tools that are based on Mutect and Mutect2 [26]. In our mutation calling analysis, we did not mark duplicates because the OS-Seq libraries are prepared without PCR amplification, i.e. a single read represents a single molecule. In addition, we masked the first 40 bases of R2 reads as N bases because they did not originate from the sample’s gDNA, but from the OS-Seq probes. We considered mutations pathogenic if they had a CADD score greater than 20 [27].

### Whole genome sequencing

To assess the accuracy of our copy number analysis, we sequenced matching tumor and normal samples from four patients using the Illumina MiSeq or NextSeq platform (Illumina, San Diego, CA, USA) **(Additional File 2: Table S5)**. Sequencing libraries were prepared using 50 ng of DNA with the KAPA HyperPlus Kit as per the manufacturer’s protocol (Roche). The libraries were sequenced with a paired-end read length of 150bp. The sequence data was automatically index-assigned (i.e. normal and tumor) and aligned with BWA [28], using default parameters, against human genome build 37. Duplicate reads were removed. The data was sorted and indexed with samtools [29]. We used the program CNVkit [30] to identify copy number alterations.

We used linked read sequencing to retain long-range genomic information from three tumor/normal sample pairs **(Additional File 2: Table S5)**. We prepared the sequencing libraries for the samples using the Chromium Library Kit (10X Genomics) following the manufacturer’s protocol. The library was sequenced using the Illumina NovaSeq 6000 system with 150 by 150-bp paired-end reads. The resulting BCL files were converted to fastq files using Long Ranger (v2.1.2) ‘mkfastq’, then Long Ranger (v2.1.2) ‘wgs’ was run to align the reads to GRCh37.1 and detect rearrangements. We called somatic variants using the Sentieon TNseq package (v201808.03) and identified copy number alterations using CNVkit [30] following the best practice guidelines.

### Digital PCR validation of copy number alterations

The digital PCR assays were run on a QX200 droplet digital PCR system (Bio-Rad) per the manufacturer’s guidelines. Additional details including the primers and thermocycling programs are described in **Table S6** and **Supplementary Methods**(see **Additional File 1**). We assessed each patient sample with three independent replicates for gene copy number. For samples analyzed in the hydrolysis probe-based assay, droplets were clustered using QuantaSoft (version1.2.10.0).

### Identification of chromosome arm alterations in TCGA CRC samples

TCGA CRC copy number alterations were downloaded [31]. CNTools R package was then used to convert the segment data to gene level data. We considered a log2 ratio greater than 0.2 as copy number gain, and less than −0.2 as copy number loss. A chromosome arm-wide event was defined as copy number alterations among 50% or more of genes located in a given chromosome arm and having a consistent gain or loss. For samples labeled as MSS, MSI-low **(MSI-L)**, or MSI-H, the frequency of arm-wide events was calculated by dividing the number of altered samples by the total number of samples in the category.

## RESULTS

### Evaluating diverse microsatellite classes and cancer drivers

We developed a cancer sequencing approach to identify somatic alterations for the major classes of tandem repeat motifs and to detect the presence of other genomic instability features, such as copy number alterations **(Figure 1a)**. In addition to mutations and copy number, this method enables simultaneous characterization of multiple features of genomic instability such as MSI, EMAST, CIN and clonal architecture. Our sequencing approach had a number of features that are significantly different than conventional targeting methods, such as bait hybridization capture or PCR amplicons [21, 22]. Specifically, this cancer sequencing technology uses multiplexed pools of primer probes that anneal to a genomic region-of-interest. One probe is used for a single target. After target-specific hybridization has occurred, the individual primer mediates a polymerase extension, replicating the downstream target DNA for sequencing **(Additional File 1: Figure S1)**. Targeting can be massively multiplexed for tens of thousands of targets if not higher. This method has several advantages. First, sequencing a specific microsatellite requires only identifying a single primer with a length in the tens rather having two select primers in PCR or using a bait hybridization oligonucleotide. This is a critical feature as many important microsatellites are flanked by repetitive sequences which complicate targeting. Second, the primer used in targeting is sequenced in addition to the adjacent genome sequence and this information provides an index specific to the microsatellite. We use this feature to improve identification and subsequent microsatellite length calling. Third, the assay can be configured to target multiple tens of thousands of targets in a single multiplexed reaction, which enables us to ascertain the status of many microsatellites in parallel. Fourth, as we show, the assessment of copy number changes is highly accurate. Finally, this approach enables us to eliminate one of the major steps in library amplification, reducing the extent of amplification errors. Given these advantages, we determined microsatellite genotypes across different motifs with high sensitivity and specificity, quantitated MSI based on this expanded signature and provided highly accurate copy number levels. We confirmed the accuracy of our copy number levels by corroborating with digital PCR and WGS.

To improve the detection of somatic alterations that occur at lower allelic fractions, we used ultra-high sequencing coverage of microsatellite and gene targets, averaging 2,865 X coverage per sample **(Additional File 2: Table S7)**. To reduce PCR artifacts, we eliminated the PCR amplification steps for library preparation **(Additional File 1: Figure S1)**. As a result of this optimization, the number of PCR-related artifacts related to amplification stutter in repetitive sequences are reduced significantly [23]. As we demonstrated previously, the elimination of PCR amplification means that each sequence read represents an individual DNA molecule [23]. Thus, there is no need to remove duplicate sequences arising from the amplification of the same molecule [23]. Removing PCR amplification improves the detection of i) microsatellite alleles which are present at low allelic fractions [23], and ii) microsatellite allele shifts which are less than 3bp in size.

We examined 225 microsatellite markers and 1,387 exons from 85 cancer genes involved in CRC biology **(Additional File 2: Table S1)**. The microsatellite markers included the following: 144 mononucleotides; 37 dinucleotides; 6 trinucleotides; 38 tetranucleotide motifs. Many of these markers had been previously used in other genotyping studies **(Additional File 2: Table S2)**. This diversity of microsatellites allowed us to distinguish different types of MSI including conventional MSI-H and EMAST. Among the microsatellite markers with mononucleotide repeats, their motif count (i.e. number of repeats) ranged from seven to 49. For the dinucleotide repeats, the motif count ranged from four to 21. For the trinucleotide repeats, the motif count ranged from nine to 18. For the tetranucleotide repeats, the motif count ranged from seven to 29. We included all five microsatellites that are part of the Bethesda criteria which includes mononucleotide repeats (BAT25, BAT26) and dinucleotide repeats (D2S123, D5S346, D17S250) [3].

Leveraging the results of TCGA, we identified 85 cancer genes that have among the highest frequency of mutations in CRC and are known cancer drivers [1]. These 85 genes were located across all autosomal chromosomes, as well as the X chromosome, and included *APC*, *TP53*, *KRAS* and other well-established cancer genes [1, 32].

### Analysis of colorectal cancer

Forty-six CRCs were used for this study **(Additional File 2: Table S8)**. A subset of the samples had prior clinical testing for the presence of MSI. All of these samples had matched tumor and normal pairs. Patients covered a range of all clinical stages: 9 had Stage I, 12 had Stage II, 8 had Stage III and 16 had Stage IV. The average age of diagnosis among the patients was 59.2 years, ranging from 31 to 83 years.

We identified somatic alterations in microsatellites, driver mutations and copy number alterations from 225 microsatellite markers and 1,387 exons from 85 cancer driver genes. We developed a new analytical method for determining microsatellite mutations and MSI quantitation. For a given microsatellite locus, we calculated the distance between two samples using all the observed microsatellite alleles **(Methods** and **Additional File 1: Figure S2a)**; we determined an allele coverage proportion vector for any given microsatellite. This algorithm leveraged the improvements in sequencing data quality that resulted from reducing amplification stutter and eliminating artifactual microsatellite alleles **(Additional File 1: Figure S3)** [23].

Next, we compared our sequencing-based microsatellite genotypes to the results from two PCR assays with different microsatellite panels and measured via CE. The first PCR assay tested microsatellites with mononucleotide repeats. The second assay tested those with tetranucleotide repeats that have been used to characterize EMAST. All of the samples were genotyped with both PCR assays. Overall, there was a high correlation between the targeted sequencing and PCR-based genotypes (R^2^ = 0.95 for mononucleotide repeats, R^2^ = 0.76 for tetranucleotide repeats; **Figure 2a**). This result indicates that the sequencing-based microsatellite genotypes were accurate.

**Figure 2.**
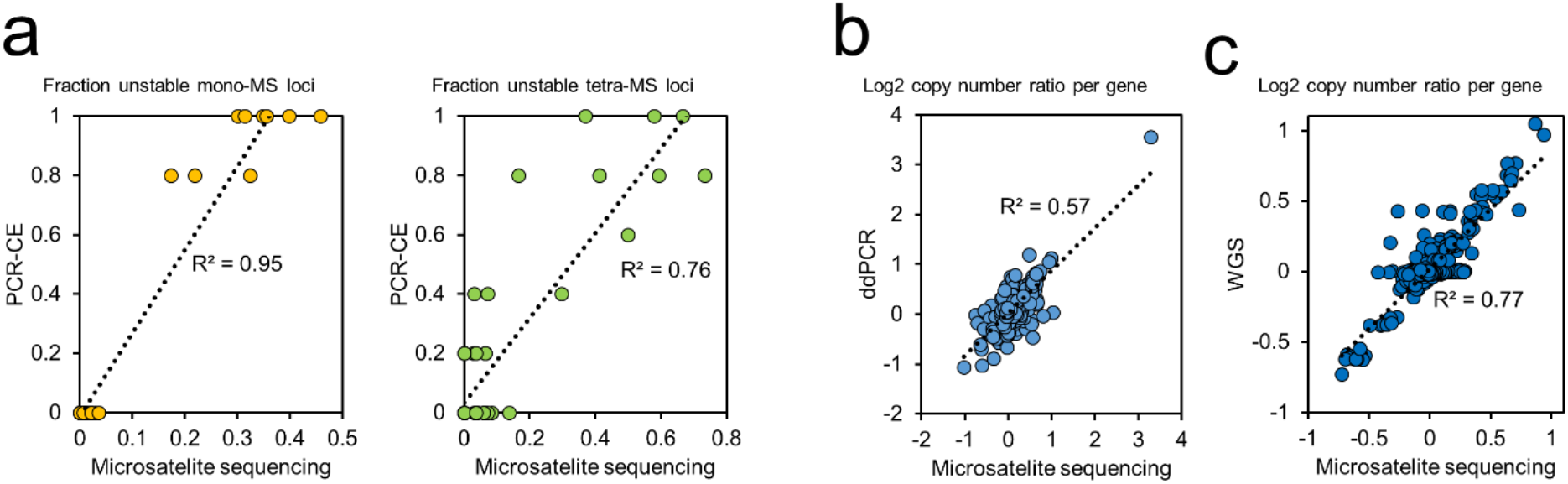
Comparison with conventional methods. **(a)** Comparison with PCR genotyping for mono- or tetranucleotide repeat microsatellites. For each of 46 CRCs, the fraction of unstable loci was calculated in each microsatellite class by dividing the number of somatic allele shift mutations by the total number of genotyped microsatellites. For PCR genotyping assays, five microsatellite markers were used for each. For the sequencing assay, 144 mono- and 38 tetranucleotide repeat microsatellites were used, respectively. **(b)** Comparison with digital PCR **(dPCR)** and whole genome sequencing **(WGS)** for gene copy numbers. For each of 46 CRCs, the log2 gene copy number ratio between tumor versus matched normal samples was calculated as described in Methods. To validate the NGS results, seven genes (*VEGFA*, *MET*, *FGFR1*, *CDK4*, *FLT3*, *ERBB2*, and *AURKA*) were selected for dPCR testing. For comparison with WGS, 83 target genes were used. In all the plots, dotted black lines indicate linear regression, and the correlation is indicated as an R-squared value.

### Calling somatic copy number alterations

To detect copy number events that co-occur with MSI, we developed a new method to accurately measure copy number alterations **(Methods; Figure 1a)**. This feature leveraged the highly reproducible targeting performance of this approach [21]. We observed that for any given DNA sample, the number of sequencing reads generated from an individual probe was highly reproducible across replicates and different DNA samples **(Additional File 1: Figures S4a and S4b)**. Primers targeting the same genomic location had the same, consistent read count. In addition, we observed that the ratio between read counts from the primers targeting the amplified or deleted regions will be different from other regions having no such changes **(Additional File 1: Figure S4c)**. To determine copy number alterations, we normalized the read counts for systematic biases associated with GC content and the read counts. We eliminated those primers with greater variance in normalized count and thus performed with lower reproducibility. Subsequently, we determined the read count ratio for each target gene and determined the ratio comparing the tumor to the matched normal genome **(Methods)**. Thus, a copy number was determined for each gene.

To validate the accuracy of our copy number measurements, we compared our sequencing-based results to other methods. We tested a subset of samples with digital PCR copy number assays for *AURKA*, *CDK4*, *FLT3*, *VEGFA*, *ERBB2*, *FGFR1* and *MET* **(Figure 2b)**. This comparison showed a high correlation between the MSI sequencing assay and digital PCR (R^2^ = 0.59), supporting the accuracy of our approach in copy number determination. In addition, we conducted WGS studies of seven sample pairs **(Additional File 2: Table S5)**, and the targeted gene copy number changes had a high correlation with the WGS results (R^2^ = 0.77; **Figure 2c**).

### Profiling CRC microsatellite instability across different sequence repeat motifs

We determined the extent of MSI across different categories of tandem repeats **(Additional File 2: Table S9)**. Among the 225 microsatellites that were sequenced, 129 of them had a somatic mutation leading to an allelic shift as detected in at least one CRC **(Additional File 2: Table S10)**. A subset did not have any somatic mutations or allelic shifts; they were characterized by short repeat lengths and included mono- and dinucleotide repeats up to 10 bp in length. In other words, all mono- and dinucleotide repeats longer than 10 bp had a mutation among the 46 CRCs. Nearly all of the other tri- and tetranucleotide repeat microsatellites (N = 44) had at least one microsatellite mutation across the entire set of CRCs. The only exception involved the tetranucleotide D4S2364, which had no mutations. Interestingly, the microsatellite mutation fractions in mono- and tetranucleotide repeats demonstrated a high correlation (R^2^ = 0.90; **Additional File 1: Figure S5**), meaning that mutations in mono- versus tetranucleotide repeats were associated.

Next, we conducted an unsupervised analysis based on our microsatellite mutation profiles. We compared our classification to established molecular testing methods. As shown in **Figure 3a**, the hierarchical clustering extrapolated from microsatellite **(MS)** mutations identified two major groups: **MS Cluster 1** (N=9) and **MS Cluster 2** (N=37). All of the CRC samples in the MS Cluster 1 had a higher percentage of unstable microsatellite loci across all the classes of microsatellites than MS Cluster 2 **(Figure 3a; Additional File 1: Figure S6)**. Overall, the mean fraction of unstable microsatellite loci was 29.0% for Cluster 1 versus 1.2% for Cluster 2 which was statistically significant (p < 0.001).

**Figure 3.**
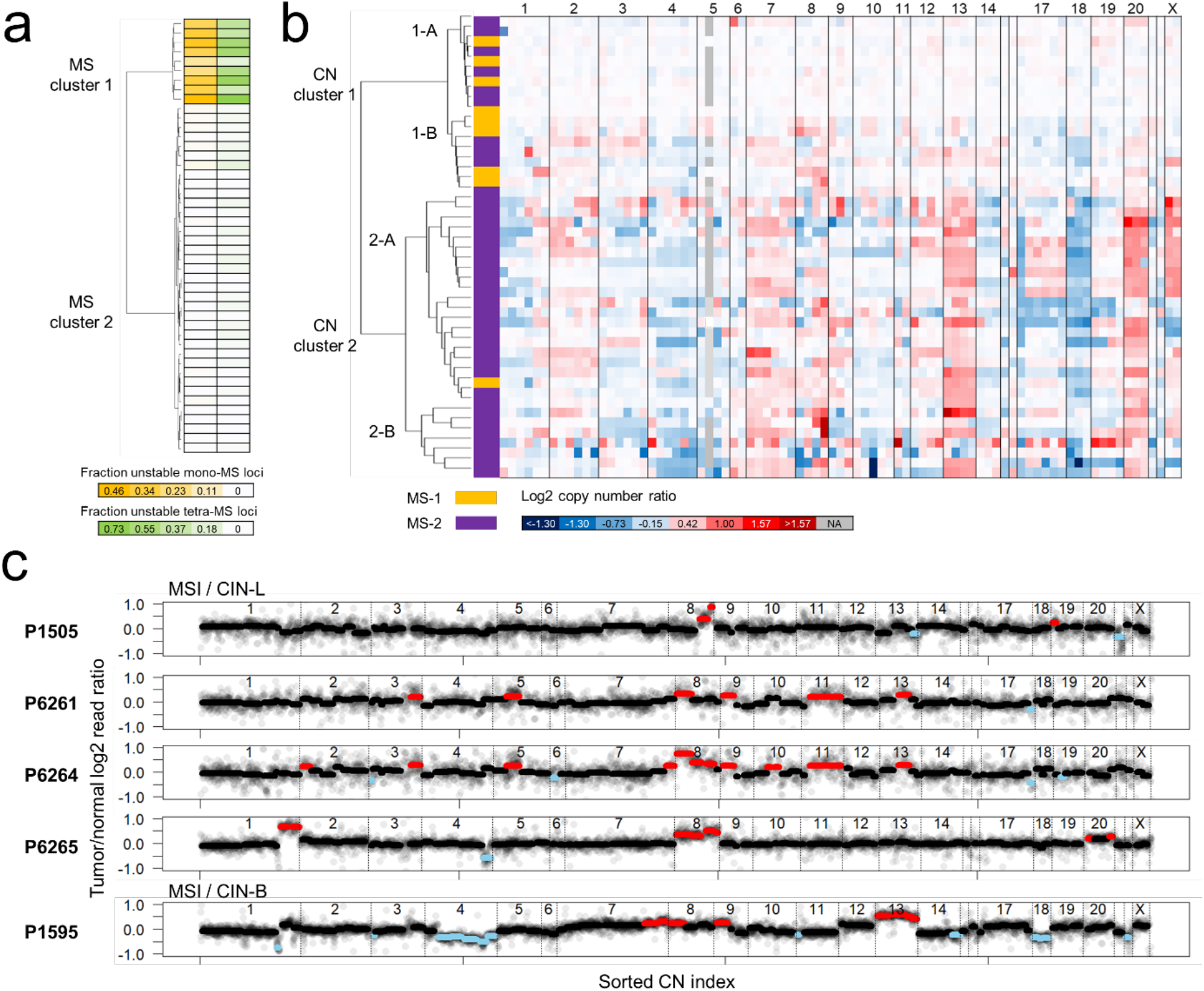
Profiling diverse sequence tandem repeats and gene copy numbers in 46 colorectal cancers. **(a)** Clustering based on 225 microsatellites across four different classes. A 225×46 matrix including the presence (1) or absence (0) of microsatellite allele shift mutations was used for an unsupervised hierarchical clustering, which generated two clusters (MS Clusters 1 and 2). The heat map of the two microsatellite classes (mono- and tetranucleotide repeats) with the most contributions are shown in two separate columns. The mutation fraction in each class was calculated by dividing the number of somatic microsatellite mutations by the total number of genotyped microsatellites. **(b)** Clustering based on tumor / normal copy number ratio of 83 target genes. The median log2 ratios for all the target genes were used for an unsupervised hierarchical clustering, which generated two major clusters. Each major cluster has two sub-clusters. In the first column of the heat map, the MS Cluster identification for each CRC is indicated as different colors. The numbers on top of the heat map indicate the chromosome where the genes are located. Amplifications and deletions are indicated with red and blue colors, respectively. **(c)** Log2 copy number ratio plots for all the CRCs having both MSI and CIN. For each CN index (x-axis), the log2 copy number ratio between read counts from tumor and normal samples (y-axis) is plotted. The median ratio value is indicated with lines of black, red, or sky blue, representing no copy number change, amplification, or deletion, respectively. The integrated MSI and CIN status is indicated on top of the plots, and the patient identification at the left of plots.

Among MS Cluster 1, all nine samples had an elevated frequency of mutations present from 17.3 to 45.9% among the different mononucleotide repeats per each tumor. In addition, all of the Cluster 1 tumors had high frequency of mutations across the different tetranucleotide repeat mutations, ranging from 16.7 to 73.3% per tumor. Our analysis included 38 tetranucleotide repeats which is higher than any other reported study. Therefore, our results showed extensive association of mononucleotide and tetranucleotide repeat instability in CRCs. In summary, MS Cluster 1 CRCs had all the features indicative of both MSI and EMAST.

Compared to MS Cluster 1, mutations among the different classes of microsatellites was significantly lower in MS Cluster 2. In MS Cluster 2, 25 out of 37 samples had mononucleotide repeat mutations. They ranged anywhere from 0.7 to 3.5% of this class of microsatellites per a tumor. Moreover, 22 out of 37 CRCs had low levels of tetranucleotide repeat instability. For MS Cluster 2, we observed a range of 2.9 to 13.8% tetranucleotide repeats with mutations per tumor.

Among the tetranucleotide microsatellites, we identified some markers that were superior to the others in terms of sensitivity or specificity between the two MS clusters **(Additional File 2: Table S10)**. Among the tumors in MS Cluster 1, the tetranucleotide marker with the highest proportion of mutations was D17S1301 (AGAT repeat) which is located at the locus 17q25.1. Seven tetranucleotide markers (F13B, THO1, TPOX, D3S3053, D5S818, D3S1358, and D14S1434) had a perfect specificity to MS Cluster 1 although the sensitivity was lower than the D17S1301 marker. These results suggest that these markers should be included when evaluating EMAST status.

### Comparison with conventional detection for MSI-H and EMAST

From the same CRC samples, we used PCR and IHC assays for a conventional determination of MSI-H and EMAST status **(Methods)**. We used an MSI PCR panel (NR-21, NR-24, BAT25, BAT26, and MONO-27) – the Bethesda criteria defining MSI-H status requires two or more microsatellites to show size shifts based on somatic alterations. CRCs with only one microsatellite with an allelic shift indicates MSI-L. Nine CRCs were MSI-H according to the PCR testing results **(Additional File 2: Table S11)**. Eight of these CRCs from MS Cluster 1 had also undergone clinical IHC testing for the four MMR proteins (MLH1, MSH2, MSH6, and PMS2) **(Additional File 2: Table S8)**. All eight of these samples lacked MMR protein expression, which was consistent with the MSI status determined from our sequencing results. Then, we tested a set of five tetranucleotide repeats (D20S82, D20S85, D8S321, D9S242, and MYCL1) previously used to determine EMAST status [12]. If two or more show a size shift, this indicates EMAST instability. Eleven CRCs were classified as EMAST-positive **(Additional File 2: Table S12)**. Finally, no tumors had evidence of MSI-L per PCR testing.

We compared the PCR genotyping results to our sequencing analysis. All nine tumors in MS Cluster 1 (sequencing) were positive for both MSI-H and EMAST PCR test. Overall, this result confirmed that our sequencing method was fully concordant with MSI-PCR. Thus, MS Cluster 1 designated tumors with both MSI-H and EMAST. Again, this validation result suggests a general association of MSI-H and EMAST.

All of the tumors in MS cluster 2 were negative for MSI-H per PCR testing **(Additional File 2: Table S11)**. This result was generally corroborated by the sequencing results where only a small fraction of mononucleotide markers had somatic mutations. In addition, we identified only a small number of CRCs that were EMAST positive via PCR. Two CRCs (P544 and P685) were EMAST positive as denoted by somatic allelic shifts in two markers based on PCR analysis with CE. However, when we compared these results to our sequencing analysis, both of the tumors showed relatively low levels of tetranucleotide repeat instability **(Additional File 2: Table S9)**. Per our sequencing analysis, the P544 CRC had 7.1% of the tetranucleotide markers with mutations. The P685 tumor had an even lower frequency at 3.1%. Given that other studies have reported a much higher frequency of EMAST among CRCs, our results suggest that a greater number of tetranucleotide markers may be required to achieve sensitive and specific identification of EMAST among these tumors. We used 38 markers for this study. Another potential implication of these results is that EMAST may represent a mixed genomic instability state in CRC. We present results supporting this possibility later.

Based on comparing NGS to PCR testing, we concluded that MS Clusters 1 and 2 were indicative of MSI and MSS status, respectively. Among this set of samples, none of the CRCs had MSI-L. In addition, we did not identify any CRCs that were exclusive to EMAST or MSI-H. Therefore, using this highly sensitive deep sequencing approach, our result show that MSI globally affects all classes of microsatellites.

### The 3-bp shift criterion improves specificity of MSI classification

We determined that microsatellite markers varied in their specificity for detecting MSI. From the sequencing results, we examined reads covering the microsatellite markers (NR-21, BAT25, BAT26, D2S123, D5S346 and D17S250) used in the Bethesda panel and three overlapping markers provided in a commercial set (Promega) **(Table 1)**. Our sequencing analysis detects any size of somatic indel shift compared to the matched normal tissue genotype, even as small as 1-bp. A 3-bp shift cutoff minimizes false positive detection due to PCR assay variation [33]. If we applied the 3-bp shift criterion to our sequencing results from the tumor, the Bethesda panel was as specific as the commercial assay in determining MSI. A previous report showed that dinucleotide markers included in the Bethesda panel were less specific for detecting cancer MSI [34]. However, our results showed that the specificity of MSI detection improved with the 3-bp shift criterion when applied to dinucleotides markers.

**Table 1.** Frequency of allele shifts in traditional MSI markers.

| STR ID | Panel | Motif | Frequency of all shifts |  | Frequency of ≥3bp shifts |  |
| --- | --- | --- | --- | --- | --- | --- |
|  |  |  | MSI<br>(N = 9) | MSS<br>(N = 37) | MSI<br>(N = 9) | MSS<br>(N = 37) |
| NR-21 | Powerplex | A | 100% | 0% | 100% | 0% |
| BAT26 | Bethesda,<br>Powerplex | A | 100% | 19% | 100% | 0% |
| BAT25 | Bethesda,<br>Powerplex | T | 100% | 28% | 100% | 0% |
| D17S250 | Bethesda | GT | 100% | 29% | 67% | 0% |
| D5S346 | Bethesda | TG | 78% | 0% | 22% | 0% |
| D2S123 | Bethesda | CA | 67% | 0% | 56% | 0% |

Using our sequencing-based results, we evaluated somatic indels of any size including 1- and 2-bp shifts. We noticed differences among the Bethesda markers that distinguish MSI. When considering somatic indels of any size, mutations in the D2S123 and D5S346 microsatellites occurred exclusively in the MS Cluster 1 tumors **(Table 1)**. The other three microsatellites (BAT25, BAT26 and D17S250) had indel mutations in both MS Cluster 1 (MSI/EMAST) and MS Cluster 2 (MSS) tumors. When we applied the 3-bp criterion, all five of the conventional Bethesda microsatellites identified only MSI tumors, thus increasing the specificity. However, this criterion reduced the sensitivity. For example, the D5S346 had microsatellite mutations in seven MSI tumors, but only two had a shift of 3 bp or more in size.

Notably, indel shifts of 3 bp or greater in length occurred only in MSI tumors, which was true not only in the microsatellites in the Bethesda panel **(Additional File 2: Table S13)**, but also in all the mono- and dinucleotide microsatellites included in our assay. In other words, only 1- or 2-bp allelic shifts for mono- and dinucleotide microsatellites were detected among the MSS tumors.

Specifically, there were 27 MSS tumors with the microsatellite indel shifts smaller than 3 bp in size: 12 had at least two microsatellites that were affected; the remaining 15 had only a single affected microsatellite. These CRCs with small microsatellite allelic shifts may have a minor sub-population with an MMR deficiency, which is only detectable with a high sensitivity method, such as our ultra-deep targeted sequencing assay. We provide an example of such a minor sub-population found in the P592 CRC later with more details.

### Analysis and classification of copy number alterations

Similar to our analysis of MSI, we conducted a separate unsupervised clustering using only the copy number **(CN)** alterations from 83 targeted genes with the most reproducible copy number calling results. Our analysis identified two major CN clusters **(Figure 3b)**. **CN Cluster 1** had a total of 18 CRCs and on average only 7% of genes had a CN. **CN Cluster 2** had the remaining 28 CRCs which had a significantly higher number of CN alterations affecting 44% of the genes on average. The differences in CN between the two clusters was highly significant (p < 0.001, **Additional File 1: Figure S7a**).

CN Clusters 1 and 2 had distinct sub-clusters **(Figure 3b)**. CN Cluster 1 had two sub-clusters **(Additional File 1: Figure S7b)**. The first cluster, **CN Cluster 1-A** (N = 10), had copy number changes averaging less than 1% of the genes per sample, which is in line with a chromosomally stable **(CS)** state. The second cluster, **CN Cluster 1-B** (N = 8), had copy number changes evident in the range of 6-26% of the genes across these samples, demonstrative of a lower level of CIN. This difference between the two subclusters was highly significant (p < 0.001). Likewise, **CN Cluster 2** had two distinct subclusters. **CN Cluster 2-B** (N = 7) had high-amplitude focal gene amplifications with six or more copies per gene or the presence of homozygous deletions. **CN Cluster 2-A** (N=21) had copy number changes that involved amplifications of broader genome segments that could extend over entire chromosome arms. A focal amplification of *MYC*, as denoted by a log2 ratio greater than 2, was present in the P98 and P685 CRCs. *MYC* is a well-known oncogenic driver associated with amplifications. The tumor suppressor gene, *TP53*, was one of the most frequently deleted genes (N = 21), as has been observed among other studies [35]. In addition, we identified a series of chromosome-wide events (i.e. amplification or deletion of all the target genes across the arm of a given chromosome). For example, chromosome-wide amplification of Chromosome 13 (N = 16), Chromosome 20 (N = 17) and deletion of Chromosome 18 (N=21) were the most frequent among the all of the CRCs.

### Determining the classes of chromosomal instability

To determine if our hierarchical clustering analysis was related to different categories of CIN, we used a multi-class statistical model based on the copy number alterations for the targeted 83 cancer genes. For training, we used copy number information from the TCGA CRC copy number dataset (N = 339) using the same genes from our panel. A previous study from Liu et al. of colorectal cancer identified specific classes of CIN involving either focal **(CIN-F)** versus broad **(CIN-B)** genomic copy number changes [36]. In this report, the size and amplitude of events were summarized by a score. A statistical threshold was used to define the two major classes. CIN-F was characterized by high amplitude focal amplifications whereas CIN-B had low-amplitude amplifications that spanned broader segments of the genome. This latter category included tumors where copy alterations covered entire chromosome arms.

For our study, we trained our 83 gene classifier using TCGA CRC results from the study of Liu et al. [36]. Then, we determined how our hierarchical CIN clustering (Cluster 1 and 2) overlapped with the CIN states (i.e. CS, CIN-F, and CIN-B) defined by Liu et al. The sensitivity and accuracy of the model was evaluated by performing five-fold cross-validation. The model had an overall prediction accuracy of 100% (sensitivity 1, specificity 1), indicating that our CIN cluster results overlapped precisely with the CIN-B and CIN-F states based on the TCGA data set. Additionally, when we reversed our training data and validation TCGA datasets, we found the same level of sensitivity and specificity. Thus, we concluded that CN Cluster 2A indicated CIN-B while CN Cluster 2B indicated CIN-F.

In terms of CIN Cluster 1, we observed a distinct subset. **Cluster 1-B** had a significantly higher number of affected genes than **Cluster 1-A (Additional File 1: Figure S7b)**. In addition, the amplitude of copy number changes in Cluster 1-B was significantly higher than that in Cluster 1-A; the difference was measured by comparing the variances of log2 gene copy number ratios (p < 0.001, **Additional File 1: Figure S7c**). Given this significant difference, we classified CRCs in CN Cluster 1-B as chromosome instability low **(CIN-L)**, an indicator of the low degree of copy number changes. The remaining CRCs were considered to be CS.

### MSI and CIN co-occurrence in CRCs

Having established that our sequencing-based profiling of microsatellites and cancer genes accurately detected various MSI and CIN classes, we determined whether there was evidence of co-occurrence of these genomic instability states. Among the nine CRCs with MSI, five had evidence of co-occurring copy number alterations and thus, indicators of CIN. Four CRCs (P1505, P6261, P6264, and P6265) had both MSI and CIN-L **(Figure 3c)**. Interestingly, all four tumors had amplifications among four genes (*WRN*, *FGFR1*, *TRPS1* and *MYC*) which are located on Chromosome 8. No deletions were noted among these genes.

In contrast, MSS CRCs had both amplifications and deletions among these same Chromosome 8 genes. Specifically, there were 31 MSS tumors with CIN-L, CIN-B or CIN-F. Nineteen of these tumors had at least one deletion among the four genes on Chromosome 8. Notably, the majority of the deletions included genes on the 8p arm; fourteen of the MSS tumors had no deletions of the genes located on the 8q arm, but only of the genes on the 8p arm.

One tumor had a striking and distinct pattern of mixed genomic instability. The P1595 CRC was MSI / CIN-B. This tumor had the highest number of chromosome-wide copy number changes among the MSI CRCs **(Figure 3c)**. We detected amplifications among the genes in Chromosome 8 and 13 as well as deletions in Chromosomes 4 and 18. This pattern matched that of the CRCs which were MSS and CIN positive. This combination of genomic instability features would have been missed with conventional MSI PCR testing. Very few, if any, CRCs have been examined for this combined genomic instability phenotype.

Liu et al.’s result from their TCGA CRC study was examined; there were multiple examples of tumors with MSI-H and CIN broad or focal copy number alterations [36]. Other studies have noted similar observations where 26 to 61% of MSI CRCs have copy number alterations or features of CIN **(Additional File 2: Table S14)** [37–44]. These other studies used conventional assays including karyotyping, fluorescent in situ hybridization **(FISH)**, flow cytometry for aneuploidy, array comparative genomic hybridization **(CGH)** and oligonucleotide arrays. Although these studies used different platforms for evaluating chromosome alterations, if one aggregates all of their samples (N=248), 41% (N = 101) showed mixed MSI and CIN features. Given that none of these studies used genome sequencing to define copy number, these previous reports may underestimate the prevalence of mixed instability classes. Moreover, they lacked the resolution of delineating specific genomic features which cancer NGS methods provide.

### Tumor mutations and overlap with genomic instability states

From the deep sequencing of the cancer genes, we identified mutations including substitutions and indels among well-established CRC driver genes **(Additional File 1: Figure S8; Additional File 2: Table S15)**. As one would expect, the MMR genes (e.g. *MLH1*, *MSH2, MSH3, MSH6*, and *PMS2*) had different mutation frequencies when comparing the MSI versus MSS CRCs **(Figure 4a)**. All nine of the MSI CRCs had at least one somatic mutation in MMR genes. Six of the MSI CRCs had germline mutations of MMR genes, and four of them also had a somatic mutation at the gene with a germline hit. The only MSS tumor with an MMR mutation was P592, which had a somatic mutation in *MSH6* as we describe in more detail later.

**Figure 4.**
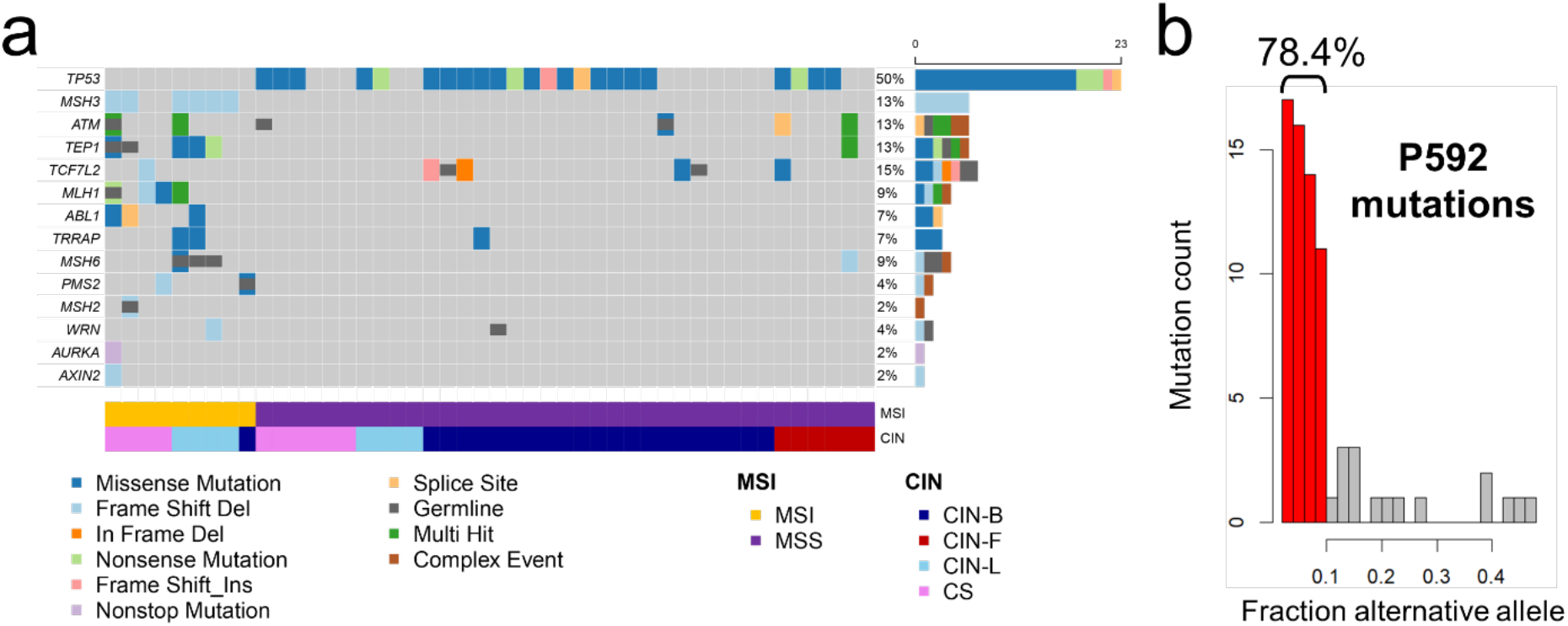
Mutation profile for genes related to DNA maintenance. **(a)** Oncoplot for the DNA maintenance genes. For all of the genes related to DNA maintenance (rows), mutation profiles of 46 tumors (columns) are shown. Only the mutations with a CADD score greater than 20 are used. Different types of somatic mutations are shown as rectangles with different colors. Gray color indicates that there is no mutation call at a given gene. Germline mutations are also indicated with a shorter rectangle overlaid on the somatic mutation map. The right panel shows the number of affected samples for each gene. Genes are sorted according to the frequency of somatic mutations. Lower panel indicates MSI and CIN sample annotations determined by the sequencing assay. **(b)** Distribution of alternative allelic fractions in the P592 tumor. The mutations with an allelic fraction less than 0.1 are indicated with red color, and the percentage is also provided on top of the corresponding histogram bars.

We examined the 16 genes (*MSH2, MSH3, MSH6, MLH1, PMS2, ATM, TP53, WRN, TRRAP, AURKA, TEP1,* etc.) which play a role in DNA repair and genome stability (**Additional File 2: Table S3**). Our results included the following: 10 genes had frequent mutations among the MSI tumors; three genes had mutations in both MSI and MSS tumors, and one gene had a mutation among the MSS tumors. Notably, MSI tumors had no mutations in *TP53* versus 62.2% of MSS CRCs that had *TP53* mutations.

An *MSH3* indel was found to be a hotspot mutation among the MSI tumors (66.7% in MSI versus none in MSS). This recurrent indel was at an adenine mononucleotide repeat, located at exon 7 of *MSH3*; this mononucleotide homopolymer is described as being eight bases in the genome reference). Interestingly, *MSH3* ‘s mutation allelic fraction had a general correlation with the extent of MSI observed in tetranucleotide repeats **(Additional File 1: Figure S9)**. Among the five CRCs with MSI and CIN, *MSH3* mutations were found in four tumors. As noted, when considering those CRCs with an *MSH3* mutation, the allele fraction of the *MSH3* mutation had a positive correlation with the degree of EMAST **(Additional File 1: Figure S9)**. *MSH3* mutations did not co-occur with *TP53* mutations. A similar level of exclusivity was evident among other sets of CRCs including those analyzed in the TCGA study where among 323 CRCs with a mutation in either gene, 95% were exclusive to one or the other [31].

A notable example of mixed genomic instability states was evident in the P592 tumor. This CRC was MSS per our sequencing analysis and the MSI PCR test. However, this CRC had 74 mutations, which was even higher than the average mutation count of MSI CRCs (43 mutations) that are normally hypermutable **(Additional File 2: Table S16)**. We observed that 78.4% of P592 CRC’s mutations occurred at a lower allelic fraction of 10% or less. This lower mutation allele fraction represented a subclonal population of tumor cells **(Figure 4b)**. Interestingly, we discovered a mutation in *MSH6* that was present at a somatic allelic faction of 6.3% and led to a frameshift. Among all of the MSS tumors, this CRC was the only one with a mutation in the MMR genes. We also noted a copy number loss per a log2 ratio of −0.11, a lower value that we attribute to the deletion being in a small proportion of tumor cells **(Additional File 4)**. Moreover, this was one of the MSS tumors that had many small somatic shifts (i.e. 1 or 2 bp) in mono- and dinucleotide microsatellites as determined through deep sequencing **(Additional File 2: Table S13)**. We detected these lower fraction mutations in this tumor because of the very high sequencing coverage (3,366X) of the target cancer genes and microsatellites. Taking all of these observations into account, these results are consistent with biallelic somatic events in *MSH6* that occurred in a subclonal population. We posit that the tumor cells in this subclone were then subject to a greater level of hypermutations consistent with MSI tumors while maintaining CIN features in parallel among the other clones. One general possibility is that the MSI-L state per the MSI PCR test may reflect a subclonal population of cells with DNA MMR loss.

### Subclonal structure of CRCs with joint MSI and CIN

We determined the subclonal structure and the relative population sizes among the ten CRCs with hypermutations **(Figure 5a; Additional File 1: Figure S10; Additional File 2: Table S17)**. Our results revealed a diverse range of tumor cellular architecture types across MSI CRCs. For this analysis, we used PyClone; this algorithm relies on targeted sequencing data with high coverage, deconvolutes the clonal architecture of tumors and estimates the subclonal cellular prevalence of somatic mutations [45].

**Figure 5.**
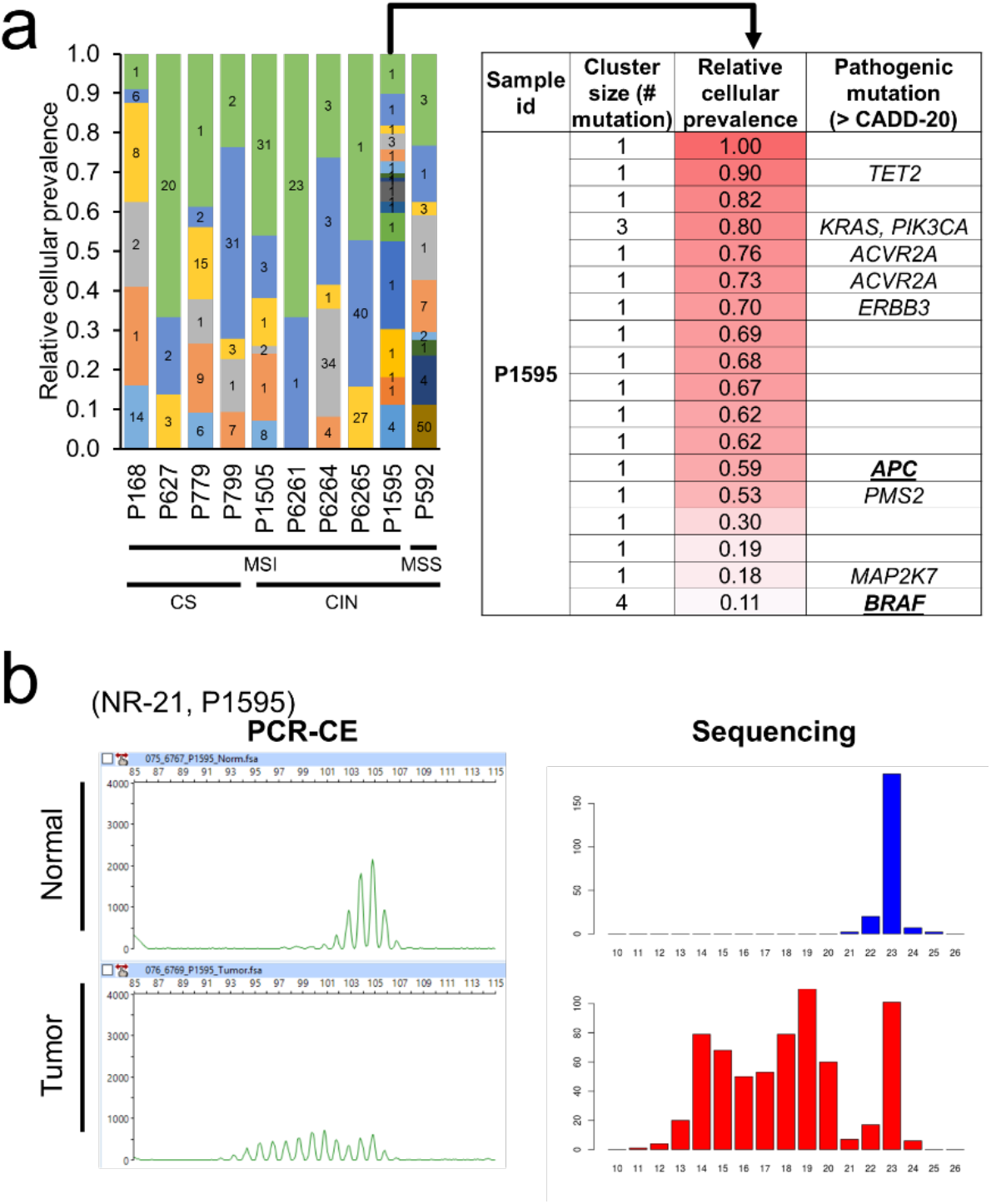
Clonal diversity of hypermutated tumors. **(a)** Clonal diversity analysis for hypermutated tumors. Relative cellular prevalence of each clone, indicated by overlapping bars in the plot, was estimated by PyClone based on allelic fraction of the mutations shared by each clonal population. The number of mutations for each clone is indicated on the inside of each bar. MSI and CIN status determined by our unsupervised analysis are indicated at the bottom. For the P592 tumor, the pathogenic mutations included in each clone are also provided in the table next to the bar plot. The underlined gene mutations (*APC* and *BRAF*) are generally known to be exclusive. **(b)** Clonal diversity shown in MSI analysis. For a microsatellite locus (NR-21), allele profiles of both normal and tumor samples from P1595 are shown. Electropherograms generated by PCR-CE method (left panels) provide relative abundance (y-axis, fluorescence intensity) of amplicons with different sizes (x-axis, DNA size in bp). Although it has stutter amplification which represents an artifact, the tumor allele profile suggests many different allele shifts (i.e. two or more), matching the observation from the mutation-based clonal diversity analysis. The allele histograms generated by our deep sequencing approach (right panels) provide relative abundance (y-axis, sequencing read count) of DNA molecules, including different microsatellite alleles (x-axis, number of motif repeats). From the normal allele profile, the homozygote allele (23 motif repeats) is apparent. On the other hand, from the tumor allele profile, many different allelic shifts are observed, also strongly supporting the clonal diversity. Given that no PCR amplification was used, these allele shifts were not results of erroneous stutter amplification but rather represented true somatic allelic shifts from mutations among different clones.

For any given hypermutated tumor, we observed different levels of subclonal diversity, which can be inferred by the number of mutation clusters. All ten CRCs had two or more clones as defined by groups/clusters of mutations with similar degrees of cellular representation **(Additional File 2: Table S17)**. Among the CRC mutations that were pathogenic (i.e. variants with a CADD score greater than 20), we noted specific patterns in terms of their clonal clustering distribution. Deleterious mutations generally occurred in larger sizes than the clusters without pathogenic variants (mean values: 1.5 versus 11.6 variants per cluster, p < 0.001) **(Additional File 1: Figure S11)**. A range of subclonal heterogeneity was observed across the MSI tumors with CIN. For example, the P6261 CRC (MSI / CIN-L) had only two mutation clusters; the cluster with largest size included 96% of the mutations.

The P1595 CRC (MSI / CIN-B) had 18 distinct mutation clusters, where the largest cluster contained only 17% of the mutations. Among the MSI tumors, this CRC had the highest level of clonal diversity as well as the highest number of copy number alterations. The high clonal diversity of P1595 CRC was also confirmed by a diverse range of tumor microsatellite mutations **(Figure 5b)**. For example, a pentanucleotide repeat marker showed six different alleles in P1595 CRC **(Additional File 1: Figure S12a)**. In contrast, the P799 CRC, with less clonal diversity, did not have as broad a range of tumor microsatellite alleles **(Additional File 1: Figure S12b)**.

### Subclonal co-occurrence of *BRAF* and *APC* driver mutations in a MSI tumor

The CRC from P1595 (MSI / CIN-B) had pathogenic mutations in both *APC* and *BRAF* **(Figure 5a)**. Somatic mutations in these two cancer drivers are generally exclusive, meaning that CRCs do not typically possess both mutations [46]. Notably, this tumor had both *APC* and the *BRAF* mutations identified in separate sub-clones based on their size. The *APC* mutation was in a mutation cluster with a cellular prevalence of 0.59, while the *BRAF* mutation was in a mutation cluster with a cellular prevalence of 0.11. Thus, we extrapolated that the *APC* mutation was a truncal cancer driver with the *BRAF* mutation appearing later in a subclonal population during the course of tumor evolution. This result suggests a dramatic divergence in this tumor where distinct genomic instability and driver pathways differentiated clonal subpopulations in an MSI tumor.

### Whole genome analysis of MSI CRCs

We applied WGS to five CRCs (P1505, P6261, P6264, P6265, and P1595) with mixed genomic instability features and two CRCs (P779 and P1710) with only one class of genomic instability **(Figure 6a; Additional File 2: Table S5)**. Specifically, the P779 CRC was MSI / CS and the P1710 CRC was MSS / CIN. To improve the detection of rearrangements, we used linked read sequencing on a subset of samples (P779, P1505 and P1595). The remainder used conventional WGS. As we have previously published, linked read sequencing uses molecular barcodes to identify and characterize single high molecular weight **(HMW)** DNA molecules up to 0.2 Mb in size, if not larger [47]. Using the long-range genomic information from those individual HMW DNA molecules, cancer rearrangements are detected with improved sensitivity, complex structural alterations are characterized more readily and Mb-scale haplotypes are revealed, some of which span entire chromosome arms [47, 48–50]. Samples had high quality HMW DNA molecules, typically in a size range of 20-40 kb on average. The HMW DNA provided phased haplotype blocks of 0.5-4.6 Mb in size **(Additional File 2: Table S5)**.

**Figure 6.**
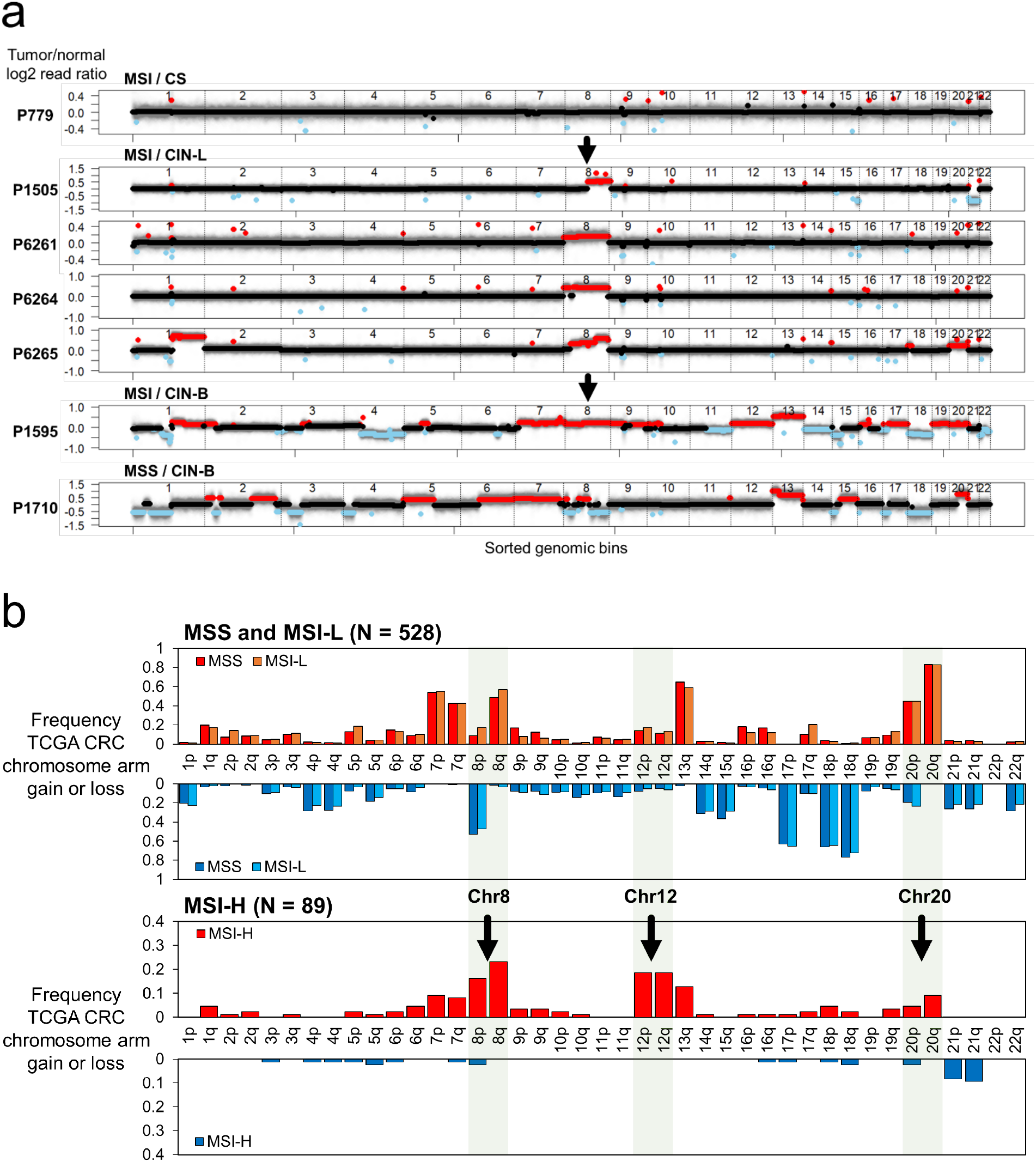
Genome-wide copy number changes in MSI tumors determined by WGS analyses. **(a)** Log2 copy number ratio plots from the WGS analysis. This analysis included all five MSI / CIN samples, as well as one MSI / CS and one MSS / CIN samples as controls. For each genomic bin (x-axis), the log2 copy number ratio calculated by CNVkit (y-axis) is plotted. The log2 ratio value of genomic segments is indicated with lines of black, red, or sky blue, representing no copy number change, amplification, or deletion, respectively. The dotted vertical lines separate genomic bins from different chromosomes. The integrated MSI and CIN status is indicated at the top of the plots, and the patient identification at the left of plots. **(b)** Validation of chromosome arm-wide copy number events in MSI tumors using TCGA CRC samples (N = 617). Separately for MSS / MSI-L and MSI-H tumors, frequencies of copy number gain and loss are shown for each chromosome arm. Gains are shown above and losses below the labels of chromosome arms.

First, we compared the WGS and targeted sequencing calls for copy number alterations **(Additional File 2: Table S18)**. We considered a genome-wide metric involving the fraction of segments with a copy number change over the entire breadth of the genome. Comparing the two results demonstrated concordance across all samples. Our targeted sequencing classification of CIN exactly matched the WGS results **(Additional File 2: Table S18)**. The CIN-B tumors (N = 2) had at least 37% or greater of their genome covered by copy number alterations. The CIN-L tumors (N = 4) had 5% or greater of their genome affected by copy number changes. Importantly, the CRCs with CIN-L (P1505, P6261, P6264, P6265) had a consistently higher degree of genomic instability than the P779 CRC with MSI / CS which was consistently diploid in its profile.

CIN events included increased copy number changes that encompassed either one or both arms of a chromosome, the latter being an example of aneuploidy. Such broad genomic copy number changes were observed in all of the MSI / CIN-L tumors – this WGS result confirmed what we observed in the targeted sequencing analysis **(Figure 6a)**. For example, the P6265 CRC (MSI / CIN-L) had a copy number increase in chromosome arms 1q, 8q and both arms of 20, indicating aneuploidy of this chromosome. P1505 had a copy number increase in chromosome arm 8q and a loss of both arms in chromosome 21. Overall, our WGS results point to the presence of notable levels of CIN in primary MSI CRCs.

### Chromosome 8 alterations among MSI colon cancers

Among the CRCs analyzed with WGS, we detected increased copy number of Chromosome 8, which corroborated our results from deep sequencing analysis. All five of the MSI CRCs with CIN features (P1505, P6261, P6264, P6265 and P1595) had an amplification of the Chr8q arm **(Figure 6a)**. The P6261 and P1595 CRCs demonstrated a copy number increase across all of Chromosome 8, indicating aneuploidy. The P6264 CRC also had Chromosome 8 aneuploidy albeit with a large segment of the Chr8p arm being diploid. P6265 had even more complex features with the telomeric segment of 8p being diploid, an increased copy number segment crossing over p and q arms and an even greater increase in the 8q telomeric segment. In contrast, the P779 tumor, classified as MSI / CS per our targeted analysis, showed no significant changes in its diploid chromosome complement.

To validate our observation among a larger number of tumors, we examined the TCGA CRCs (N = 617) and their genomic copy number data. Based on the CN profiles we determined the status of chromosome arm copy number **(Methods)**. The 8q amplification was very common among MSI-H tumors (N = 89) with 23% (N = 21) having evidence of this event **(Figure 6b)**. In addition, 16.3% of TCGA MSI CRCs had an increase in the 8p arm copy number. Likewise, MSI CRCs had frequent copy number alterations affecting Chromosomes 7, 12, 13 and 20. Regardless, the most frequent change was observed in the 8q arm.

Several other independent studies validated these findings. Using metaphase CGH, Camps et al. identified that 36% (N=5) among a set of MSI CRCs (N=13) had a chromosome 8 copy number increase [39]. Trautmann et al. used array CGH and determined that copy number increases in the locus 8q22-24 was the most commonly observed event occurring in 35% (N = 8) among a set of MSI CRCs (N = 23) [40]. With a variety of methods, other studies have also corroborated these findings involving Chromosome 8 [37, 39, 42, 43].

### Chromosome 8 translocation in an MSI CRC

With linked read WGS, we discovered an inter-chromosomal rearrangement event in the P1505 CRC (MSI / CIN-L) **(Figure 7)**. We use linked read sequence data to identify chromosome arm alterations that are assigned a specific haplotype. Cancer allelic imbalances increases the copy number representation of a specific haplotype. Using linked read sequencing from normal tumor pairs, we can determine if a chromosome arm has been involved in a duplication event that increases the ratio of one haplotype to another [48]. Referred to as digital karyotyping, this analysis produces information similar to conventional karyotyping or FISH but also has the advantage of having the resolution of whole genome sequencing. After conducting this haplotype analysis, it became clear that a specific haplotype of Chr 8q had been duplicated beginning at a specific breakpoint in the q arm proximal to the centromere **(Figure 7a).**

**Figure 7.**
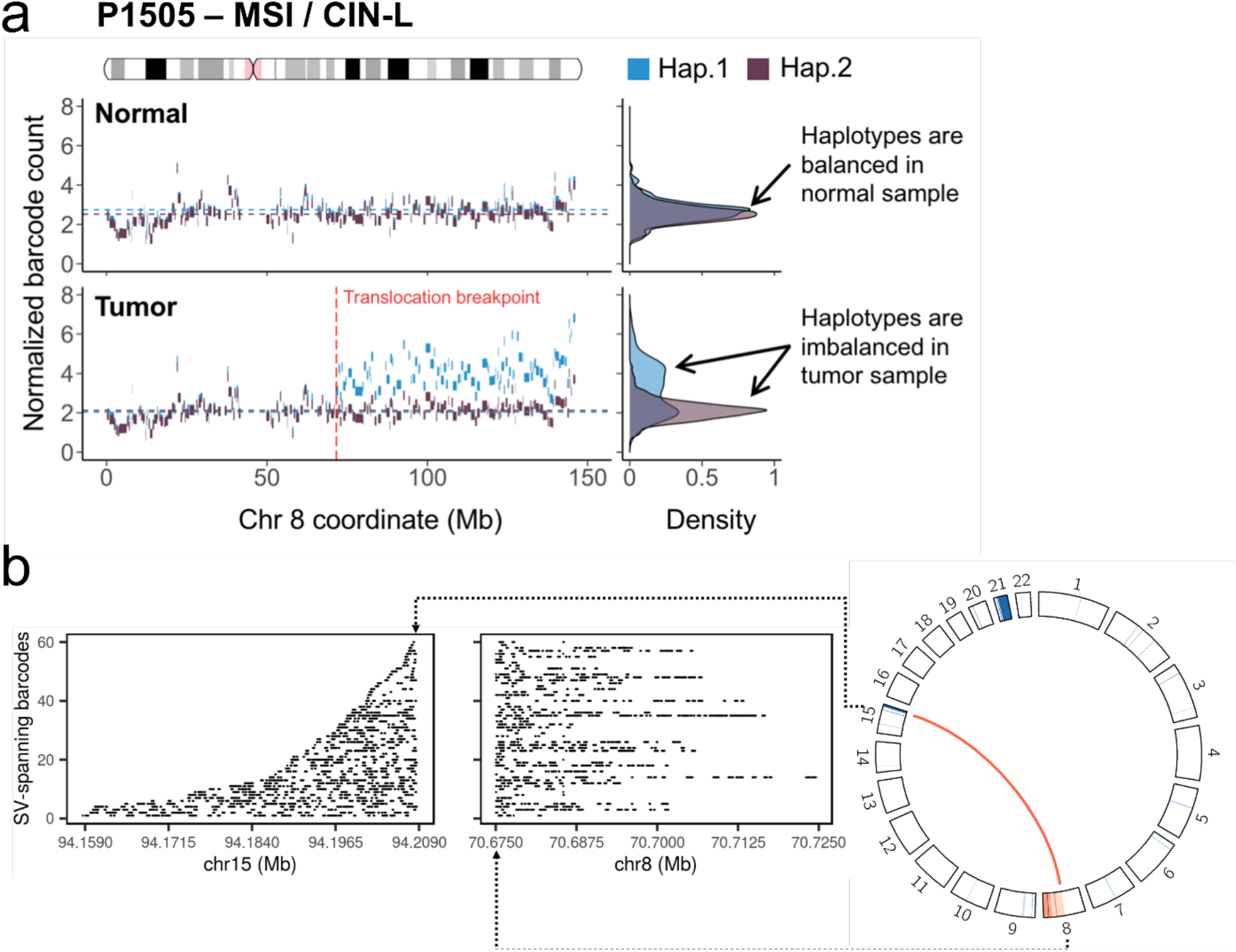
Inter-chromosomal rearrangement in a CRC tumor with a mixed MSI / CIN phenotype. **(a)** Haplotypes of chromosome 8 in the normal and tumor samples of P1505. The blocks indicate the original haplotype blocks determined by linked read sequencing, and their color denotes their parental assignment. Haplotype 1 (blue) in the tumor sample displays an allelic imbalance on chromosome 8q that reflects an amplification event, beginning at the translocation breakpoint with chromosome 15. The density plots to the right reflect the distribution of the haplotype counts. **(b)** The translocation event from the tumor sample of P1505 (an MSI / CIN-L tumor). For the CIRCOS plot (right), an inter-chromosomal change is indicated with an orange line. Chromosomes are indicated as curved boxes along the circle, with chromosome 1 at the 12 o’clock position, and continuing in a clockwise direction. The width of the box represents the size of chromosome. Inside the boxes, the log2 copy number ratio from genomic segments is displayed as a heat map, where orange and blue colors indicate amplification and deletion, respectively. The positions where an amplification at Chromosome 8 and a deletion at Chromosome 15 start coincide with the translocation breakpoints. As shown in the left panels, molecular barcodes from linked read sequences are found in two genomic regions flanking the breakpoints of the translocation. Each row indicates individual DNA molecules that are found in both the genomic regions. The alignment of barcoded reads is indicated by horizontal lines located at the genomic positions (x-axis).

On closer analysis of this breakpoint, we identified a novel translocation between Chromosomes 8 and 15 that has never been reported in colon cancer **(Figure 7b)**. The breakpoints were located at 8q13.3 and 15q26.2 and, and the Chromosome 8 breakpoint separates exons 2 and 3 of *XKR9*. This gene is a cell membrane bound protein and a member of the Kell Blood Group complex subunit-related gene family [51]. Based on a National Library of Medicine Pubmed literature search during the submission of this manuscript, there were no articles describing functional studies of this gene. One recent article reported that *XKR9* gene expression in cancer was associated with decreased overall survival [51]. In this report, Li et al. used the TCGA genomic data from 21 cancer types and identified differential expressed genes in tumors originating from different ethnicities. *XKR9* differential expression had the most significant association with survival. Another report identified recurrent deletions at the 8q13.3 locus that are causative for Branchio-Oto-Renal syndrome [52]. The deletion breakpoints involved human endogenous retroviral sequences and the authors hypothesized that these repetitive elements mediated a recombination that led to recurrent deletions causative for this genetic syndrome. Overall, our result suggests that Chromosome 8 has a predilection for CIN among MSI CRCs.

P1505 CRC harbored the *MSH3* frameshift mutation with the highest allelic fraction (84.6%) among all MSI tumors. One possibility is that the *MSH3* mutation was an early mutation during cancer development or provided a fundamental selective advantage that allowed this clone to become dominant.

## DISCUSSION

We developed an ultra-deep sequencing approach for characterizing somatic alterations of microsatellites and copy number changes in cancer. We conducted an integrative analysis of genomic instability features on CRCs. Unlike nearly all studies examining microsatellites, we examined different classes of microsatellite markers for somatic mutations indicative of MS, and thus determined quantitative instability levels, not only in mono- and di-nucleotide repeats, but also in tri- and tetranucleotide markers. Tri- and tetranucleotide repeats are generally excluded from short read sequencing-based analysis; many of them are too long to be spanned by short sequence reads [7]. Moreover, our targeted assay measured copy number changes with a precision comparable to WGS. Generally, targeted approaches such as whole exome sequencing are not considered ideal for copy number analysis [53], but our approach proved to be particularly robust as we validated with a variety of other methods including digital PCR and WGS. As an added feature, given the high sequencing coverage and elimination of many of the biases of PCR amplification, we quantified the allelic fraction of mutations accurately and used this information combined with copy number to delineate subclonal features.

MSI status was assigned for tumors having instability in all of the microsatellite classes. These MSI tumors showed distinct characteristics: i) The MSI tumors had varying degrees of microsatellite mutation with both mono- and dinucleotide repeats being proportionally elevated along with tri- and tetranucleotide repeat alterations; ii) some MSI tumors showed CIN with chromosome- or chromosome arm-wide copy number changes as well as a translocation; iii) our simultaneous MSI profiling across a larger number of microsatellites and clonal architecture deconvolution revealed examples of MSI subclones coexisting with other clonal populations. This added feature of intratumoral heterogeneity may contribute to different tumor phenotypes.

Traditional PCR tests sometimes lead to a classification of ‘low’ status, generally defined as positive only in a portion of markers such as what is observed for MSI-L and EMAST-L. Like the traditional MSI tests, PCR tests for EMAST tumors also use only five or more markers [12–14, 54]. Some recent studies used a commercial assay using 16 forensic markers [15], but the expanded number led to a new designation of EMAST-L tumors, i.e. inconclusive between positive and negative. In our study, with an expanded number of tetranucleotide markers, we obtained definitive results about the extent of EMAST, its association with MSI and did not identify this EMAST-L category. Our results suggest that a wider range and greater number of tetranucleotide markers are important for accurate determination of EMAST.

Our study showed that the MSI tumors had different microsatellite mutation fractions in a range **(Additional File 2: Table S9)**. Although with a limited resolution, the MSI PCR test using five mononucleotide markers generated results matching the microsatellite mutation fraction measured by our sequencing analysis ( R^2^ = 0.95**; Figure 2a)**. A recent study about genetic heterogeneity of MSI tumors revealed that the overall genomic MSI level, termed ‘MSI intensity’ in the study, is a predictor of response to immunotherapies [55]. Given the clinical implication, an improved MSI test would go beyond a simple positive versus negative indication, but would measure the quantitative extent of MSI. The precise quantification of the genomic MSI level will optimize patient care decisions, and our microsatellite assay provides this feature.

We separated MSI markers into two groups according to their repeat motif length (mono- and di-nucleotide repeats versus tri- and tetranucleotide repeats), and then compared microsatellite mutation rates within each group. There was a clear correlation between length of microsatellites and the mutation frequency across all tumors regardless of their MSI status **(Additional File 1: Figure S13)**. This result shows that many MSI markers are not specific, especially when they are long. All the traditional microsatellite markers are relatively long (>20 bp), and some of them were frequently mutated even in MSS tumors **(Table 1 and Additional File 2: Table S10)**. To be both sensitive and specific in MSI detection, any molecular test should include more markers with enhanced specificity (e.g. markers with intermediate length).

Another noteworthy feature of MSI tumors was that the microsatellite mutation fractions in mono- and tetranucleotide repeats were highly correlated (R^2^ = 0.90; **Additional File 1: Figure S5**). Therefore, all the MSI tumors unstable in mono- and dinucleotide repeats were also unstable in tri- and tetranucleotide repeats which define the EMAST molecular phenotype. We did not identify CRCs with instability that were exclusive to mono- and dinucleotide repeats or tri- and tetranucleotide repeats. Most studies reported tumors with only a single type of instability (i.e. MSI-H only or EMAST-positive only), which were as frequent as the tumors having both types of instability [12-15, 54]. The PCR tests can generate false positives depending on the marker choices. The original Bethesda marker set (two mononucleotide and three dinucleotide repeats) were revised because the tumors having microsatellite shifts only in dinucleotide repeat loci were frequently false positives [34]. However, the dinucleotide markers are still being used when determining MSI status [14, 54]. Moreover, there is no standardized set of markers for EMAST. We used an expanded set of 38 tetranucleotide microsatellites for identifying EMAST – our data showed a clear association with mononucleotide microsatellites **(Additional File 1: Figure S5)** and identified two false positives **(Additional File 2: Table S9)**. In summary, the MSI tumors identified by PCR tests may not characterize the full diversity of microsatellite instability across different motif repeats.

The EMAST phenotype in MSI tumors has not been examined with whole genome sequencing or other sequencing methods. It is thought that a major driver of EMAST in MSS tumors is *MSH3* loss-of-function. Changes in MSH3 cellular location rather than deleterious mutations or epimutations are the basis of this change in function [11]. However, MSI-H tumors also have a propensity for acquiring pathogenic indel mutations in *MSH3* [56]; in our study, six of nine MSI tumors had this *MSH3* indel and the mutation allelic fraction had a positive correlation with the fraction of tetranucleotide repeat loci having a microsatellite allele shift **(Additional File 1: Figure S9).** This result suggests that those MSI tumors may have acquired a later *MSH3* mutation which elicits an EMAST phenotype in a subclonal population. One or two bp deletions in a specific *MSH3* homopolymer results in a frameshift in protein expression, and eventually, an increased microsatellite mutation rate in tetranucleotide repeats. In summary, our study shows that MSI-H tumors are likely to acquire the EMAST phenotype as a result of the initial instability in mononucleotide repeats. Other studies of CRC have reported *MSH3* mutations but given their reliance on lower coverage methods (in the hundreds at most) such as exome sequencing, they lack the sensitivity to detect s the hotspot indel that we identified. Indels in microsatellites are particularly challenging to identify and thus may obscure their detection from cancer genomes.

Generally, it is thought that dysfunction among different DNA repair mechanisms leads to exclusive states of genomic instability, such as MSI or CIN. Previous studies using array CGH had identified co-occurrence of MSI and copy number alterations [40, 44]. These studies did not use genome sequencing nor did they identify tetranucleotide repeat instability in relationship to different classes of CIN. In our study, the majority of MSI tumors had both EMAST and CIN features (four CIN-L and a CIN-B), indicating a mixed genomic instability state. The extent of CIN seen among these mixed state tumors were significantly higher than among chromosome stable tumors **(Additional File 1: Figures S7b and c)**. Specifically, chromosome arm changes and aneuploidy were clearly evident, especially in Chromosome 8. Interestingly, all of the MSI / CIN-L tumors had a frameshift mutation in *MSH3*, which also supported our new classification. Changes in MSH3 function may lead to double strand breaks and chromosomal rearrangements [11]. In line with MSH3’s functional role in DNA maintenance, the tumor with the highest *MSH3* allele fraction (P1505) in our study had an inter-chromosomal rearrangement event **(Figure 7)**. The only MSI tumor with CIN-B did not have the *MSH3* frameshift mutation – one possibility is that this tumor had a different aberrant pathway leading to broad CIN. Overall, these unexpected structural rearrangements in MSI tumors suggest the presence of genomic heterogeneity of CRC tumors. Therefore, to be more precise in assessing the genomic properties of MSI tumors, we would recommend that determining the CIN status should be a supplementary biomarker. Our method enables an accurate determination of both types of instability.

CIN tumors are not responsive to immune checkpoint therapies. It is possible that some MSI positive tumors may not respond to immunotherapy due to the clonal divergence and presence of subclonal tumor cell populations with CIN characteristics. We are pursuing studies to determine if mixed MSI and CIN states alter immunotherapy response.

Tumor mutation burden is a frequently used biomarker that has recently been tested for its usage in immunotherapy response prediction; approximately 50% of TMB-high tumors still do not respond to immunotherapies [18]. Recently, a study was conducted of 67,000 patient samples including one thousand MSI-positive tumors [6]; the conventional definition of MSI was sufficient, but not necessary, for classification as elevated TMB. The Cancer Genome Atlas Project’s CRC study (N = 276) also reported the same relationship between MSI and TMB [1], where 23% of hypermutated, TMB-high CRC tumors tested negative for MSI, which included the six with the highest mutation rates. A proportion of these tumors are attributable to the altered function of DNA polymerase D1 (*POLD1*) or DNA polymerase E (*POLE*) albeit this accounts for only a small fraction of CRCs [57]. We also found such a case in our samples. The P592 CRC was an MSS tumor from our targeted sequencing-based analysis, but had a TMB as high as the MSI tumors. In addition, from this tumor we identified a low-level MSI with sequencing, which were otherwise undetectable by a conventional PCR-CE tests, and a pathogenic *MSH6* mutation with a low allelic fraction. Thus, more sophisticated MSI NGS tests which consider other critical features such as CIN and clonal architecture may provide a more accurate predictor for determining patients who may respond to immunotherapy.

## CONCLUSION

Overall, we developed a new sequencing approach that determines MSI status based on all of the microsatellite classes, CIN status and subclonal features. We found that the CIN phenotype was unexpectedly common in our MSI tumors, which suggests that evaluating CIN status may prove useful for determining immunotherapy response. Other studies validated our conclusions. Chromosome 8 shows alterations in the context of MSI and CIN. In addition, the microsatellite frameshift at exon 7 of *MSH3* and the degree of EMAST were associated with the mixed phenotype. This analysis of highly multiplexed microsatellites provided better quantitative accuracy and distinguished MSI tumors with distinct characteristics in mutation patterns in comparison to MSS tumors.

## ABBREVIATIONS

CE: capillary electrophoresis
CGH: comparative genomic hybridization
CIN: chromosome instability
CIN-B: chromosome instability broad
CIN-F: chromosome instability focal
CIN-L: chromosome instability low
CN: copy number
CRC: colorectal carcinoma
CS: chromosomally stable
EMAST: elevated microsatellite alterations at selected tetranucleotide repeats
FISH: fluorescent in situ hybridization
HMW: high molecular weight
IHC: immunohistochemistry
Indel: insertions or deletions
MMR: mismatch repair
MS: microsatellites
MSI: microsatellite instability
MSI-H: microsatellite instability high
MSI-L: microsatellite instability low
MSS: microsatellite stable
OS-Seq: oligonucleotide-selective sequencing
R1: Read 1
R2: Read 2
STRs: short tandem repeats
TCGA: The Cancer Genome Atlas Project
WGS: whole genome sequencing

## DECLARATIONS

### Ethics approval and consent to participate

This study was conducted in compliance with the Helsinki Declaration. The Institutional Review Board at Stanford University School of Medicine approved the study protocols (IRB-11886 and IRB-48492).

### Consent for publication

All patients consented for publication.

### Availability of data and material

Sequence data is available at the NIH’s dbGAP repository, accession number phs001914.v1.p1.

### Competing interests

The authors declare that they have no competing interests.

### Funding

The work was supported by the Stanford-Coulter Translational Research Grant and a Stanford Cancer Institute Translational Research Grant. This work was also supported by the National Institutes of Health grants [R01HG006137-04 to HPJ, P01HG00205ESH to GS and HPJ, R33CA174575 to EH and HPJ]. Additional support to HPJ came from the Research Scholar Grant, RSG-13-297-01-TBG from the American Cancer Society and the Clayville Foundation.

### Authors’ contribution

G.S., H.L., E.H. and H.P.J. designed the experiments. G.S., E.H., L.M., and A.F.A. conducted the experiments. G.S., S.M.G., and S.U.G. developed the analysis algorithms. G.S., S.M.G., S.U.G., L.Z., C.S., S.H. and H.P.J. analyzed the data. G.S. and H.P.J. wrote the manuscript. H.P.J. supervised the overall project.

## Acknowledgements

We thank Madeline McNamara for helpful comments.

## ADDITIONAL FILES

**Additional file 1**: Supplementary Method, Supplementary Figures S1 – S13. Format: PDF

**Additional file 2:** Supplementary Tables S1 – S18. Format: XLSX

**Additional file 3:** Sequences of OS-Seq primer probes. Format: XLSX

**Additional file 4:** Log2 copy number ratio per gene from the targeted ultra-deep sequencing. Format: TXT

## Notes

### Competing Interest Statement

The authors have declared no competing interest.

## REFERENCES

1. Cancer Genome Atlas N: Comprehensive molecular characterization of human colon and rectal cancer. Nature 2012, 487: 330–337.

2. Bacher JW, Flanagan LA, Smalley RL, Nassif NA, Burgart LJ, Halberg RB, Megid WM, Thibodeau SN: Development of a fluorescent multiplex assay for detection of MSI-High tumors. Dis Markers 2004, 20: 237–250.

3. Boland CR, Thibodeau SN, Hamilton SR, Sidransky D, Eshleman JR, Burt RW, Meltzer SJ, Rodriguez-Bigas MA, Fodde R, Ranzani GN, Srivastava S: A National Cancer Institute Workshop on Microsatellite Instability for cancer detection and familial predisposition: development of international criteria for the determination of microsatellite instability in colorectal cancer. Cancer Res 1998, 58: 5248–5257.

4. Cohen R, Hain E, Buhard O, Guilloux A, Bardier A, Kaci R, Bertheau P, Renaud F, Bibeau F, Flejou JF, et al: Association of Primary Resistance to Immune Checkpoint Inhibitors in Metastatic Colorectal Cancer With Misdiagnosis of Microsatellite Instability or Mismatch Repair Deficiency Status. JAMA Oncol 2019, 5: 551–555.

5. Bartley AN, Luthra R, Saraiya DS, Urbauer DL, Broaddus RR: Identification of cancer patients with Lynch syndrome: clinically significant discordances and problems in tissue-based mismatch repair testing. Cancer Prev Res (Phila) 2012, 5: 320–327.

6. Trabucco SE, Gowen K, Maund SL, Sanford E, Fabrizio DA, Hall MJ, Yakirevich E, Gregg JP, Stephens PJ, Frampton GM, et al: A Novel Next-Generation Sequencing Approach to Detecting Microsatellite Instability and Pan-Tumor Characterization of 1000 Microsatellite Instability-High Cases in 67,000 Patient Samples. J Mol Diagn 2019.

7. Cortes-Ciriano I, Lee S, Park WY, Kim TM, Park PJ: A molecular portrait of microsatellite instability across multiple cancers. Nat Commun 2017, 8: 15180.

8. Hause RJ, Pritchard CC, Shendure J, Salipante SJ: Classification and characterization of microsatellite instability across 18 cancer types. Nat Med 2016, 22: 1342–1350.

9. Middha S, Zhang L, Nafa K, Jayakumaran G, Wong D, Kim HR, Sadowska J, Berger MF, Delair DF, Shia J, et al: Reliable Pan-Cancer Microsatellite Instability Assessment by Using Targeted Next-Generation Sequencing Data. JCO Precis Oncol 2017, 2017.

10. Waalkes A, Smith N, Penewit K, Hempelmann J, Konnick EQ, Hause RJ, Pritchard CC, Salipante SJ: Accurate Pan-Cancer Molecular Diagnosis of Microsatellite Instability by Single-Molecule Molecular Inversion Probe Capture and High-Throughput Sequencing. Clin Chem 2018, 64: 950–958.

11. Carethers JM: Microsatellite Instability Pathway and EMAST in Colorectal Cancer. Curr Colorectal Cancer Rep 2017, 13: 73–80.

12. Watson MM, Lea D, Rewcastle E, Hagland HR, Soreide K: Elevated microsatellite alterations at selected tetranucleotides in early-stage colorectal cancers with and without high-frequency microsatellite instability: same, same but different? Cancer Med 2016, 5: 1580–1587.

13. Torshizi Esfahani A, Seyedna SY, Nazemalhosseini Mojarad E, Majd A, Asadzadeh Aghdaei H: MSI-L/EMAST is a predictive biomarker for metastasis in colorectal cancer patients. J Cell Physiol 2019, 234: 13128–13136.

14. Chen MH, Chang SC, Lin PC, Yang SH, Lin CC, Lan YT, Lin HH, Lin CH, Lai JI, Liang WY, et al: Combined Microsatellite Instability and Elevated Microsatellite Alterations at Selected Tetranucleotide Repeats (EMAST) Might Be a More Promising Immune Biomarker in Colorectal Cancer. Oncologist 2019.

15. Wang Y, Vnencak-Jones CL, Cates JM, Shi C: Deciphering Elevated Microsatellite Alterations at Selected Tetra/Pentanucleotide Repeats, Microsatellite Instability, and Loss of Heterozygosity in Colorectal Cancers. J Mol Diagn 2018, 20: 366–372.

16. Koi M, Tseng-Rogenski SS, Carethers JM: Inflammation-associated microsatellite alterations: Mechanisms and significance in the prognosis of patients with colorectal cancer. World J Gastrointest Oncol 2018, 10: 1–14.

17. Ganesh K, Stadler ZK, Cercek A, Mendelsohn RB, Shia J, Segal NH, Diaz LA, Jr.: Immunotherapy in colorectal cancer: rationale, challenges and potential. Nat Rev Gastroenterol Hepatol 2019, 16: 361–375.

18. Chan TA, Yarchoan M, Jaffee E, Swanton C, Quezada SA, Stenzinger A, Peters S: Development of tumor mutation burden as an immunotherapy biomarker: utility for the oncology clinic. Ann Oncol 2019, 30: 44–56.

19. Le DT, Durham JN, Smith KN, Wang H, Bartlett BR, Aulakh LK, Lu S, Kemberling H, Wilt C, Luber BS, et al: Mismatch repair deficiency predicts response of solid tumors to PD-1 blockade. Science 2017, 357: 409–413.

20. Walk EE, Yohe SL, Beckman A, Schade A, Zutter MM, Pfeifer J, Berry AB, College of American Pathologists Personalized Health Care C: The Cancer Immunotherapy Biomarker Testing Landscape. Arch Pathol Lab Med 2020, 144: 706–724.

21. Hopmans ES, Natsoulis G, Bell JM, Grimes SM, Sieh W, Ji HP: A programmable method for massively parallel targeted sequencing. Nucleic Acids Res 2014, 42: e88.

22. Myllykangas S, Buenrostro JD, Natsoulis G, Bell JM, Ji HP: Efficient targeted resequencing of human germline and cancer genomes by oligonucleotide-selective sequencing. Nat Biotechnol 2011, 29: 1024–1027.

23. Shin G, Grimes SM, Lee H, Lau BT, Xia LC, Ji HP: CRISPR-Cas9-targeted fragmentation and selective sequencing enable massively parallel microsatellite analysis. Nat Commun 2017, 8: 14291.

24. Chalmers ZR, Connelly CF, Fabrizio D, Gay L, Ali SM, Ennis R, Schrock A, Campbell B, Shlien A, Chmielecki J, et al: Analysis of 100,000 human cancer genomes reveals the landscape of tumor mutational burden. Genome Med 2017, 9: 34.

25. Chen TQ, Guestrin C: XGBoost: A Scalable Tree Boosting System. Kdd'16: Proceedings of the 22nd Acm Sigkdd International Conference on Knowledge Discovery and Data Mining 2016: 785–794.

26. Cibulskis K, Lawrence MS, Carter SL, Sivachenko A, Jaffe D, Sougnez C, Gabriel S, Meyerson M, Lander ES, Getz G: Sensitive detection of somatic point mutations in impure and heterogeneous cancer samples. Nat Biotechnol 2013, 31: 213–219.

27. Rentzsch P, Witten D, Cooper GM, Shendure J, Kircher M: CADD: predicting the deleteriousness of variants throughout the human genome. Nucleic Acids Res 2019, 47: D886–D894.

28. Li H, Durbin R: Fast and accurate short read alignment with Burrows-Wheeler transform. Bioinformatics 2009, 25: 1754–1760.

29. Li H, Handsaker B, Wysoker A, Fennell T, Ruan J, Homer N, Marth G, Abecasis G, Durbin R, Genome Project Data Processing S: The Sequence Alignment/Map format and SAMtools. Bioinformatics 2009, 25: 2078–2079.

30. Talevich E, Shain AH, Botton T, Bastian BC: CNVkit: Genome-Wide Copy Number Detection and Visualization from Targeted DNA Sequencing. PLoS Comput Biol 2016, 12: e1004873.

31. Cerami E, Gao J, Dogrusoz U, Gross BE, Sumer SO, Aksoy BA, Jacobsen A, Byrne CJ, Heuer ML, Larsson E, et al: The cBio cancer genomics portal: an open platform for exploring multidimensional cancer genomics data. Cancer Discov 2012, 2: 401–404.

32. Lee H, Flaherty P, Ji HP: Systematic genomic identification of colorectal cancer genes delineating advanced from early clinical stage and metastasis. BMC Med Genomics 2013, 6: 54.

33. Bacher JW, Sievers CK, Albrecht DM, Grimes IC, Weiss JM, Matkowskyj KA, Agni RM, Vyazunova I, Clipson L, Storts DR, et al: Improved Detection of Microsatellite Instability in Early Colorectal Lesions. PLoS One 2015, 10: e0132727.

34. Umar A, Boland CR, Terdiman JP, Syngal S, de la Chapelle A, Ruschoff J, Fishel R, Lindor NM, Burgart LJ, Hamelin R, et al: Revised Bethesda Guidelines for hereditary nonpolyposis colorectal cancer (Lynch syndrome) and microsatellite instability. J Natl Cancer Inst 2004, 96: 261–268.

35. Xia LC, Van Hummelen P, Kubit M, Lee H, Bell JM, Grimes SM, Wood-Bouwens C, Greer SU, Barker T, Haslem DS, et al: Whole genome analysis identifies the association of TP53 genomic deletions with lower survival in Stage III colorectal cancer. Sci Rep 2020, 10: 5009.

36. Liu Y, Sethi NS, Hinoue T, Schneider BG, Cherniack AD, Sanchez-Vega F, Seoane JA, Farshidfar F, Bowlby R, Islam M, et al: Comparative Molecular Analysis of Gastrointestinal Adenocarcinomas. Cancer Cell 2018, 33: 721–735 e728.

37. Li LS, Kim NG, Kim SH, Park C, Kim H, Kang HJ, Koh KH, Kim SN, Kim WH, Kim NK, Kim H: Chromosomal imbalances in the colorectal carcinomas with microsatellite instability. Am J Pathol 2003, 163: 1429–1436.

38. Sinicrope FA, Rego RL, Halling KC, Foster N, Sargent DJ, La Plant B, French AJ, Laurie JA, Goldberg RM, Thibodeau SN, Witzig TE: Prognostic impact of microsatellite instability and DNA ploidy in human colon carcinoma patients. Gastroenterology 2006, 131: 729–737.

39. Camps J, Armengol G, del Rey J, Lozano JJ, Vauhkonen H, Prat E, Egozcue J, Sumoy L, Knuutila S, Miro R: Genome-wide differences between microsatellite stable and unstable colorectal tumors. Carcinogenesis 2006, 27: 419–428.

40. Trautmann K, Terdiman JP, French AJ, Roydasgupta R, Sein N, Kakar S, Fridlyand J, Snijders AM, Albertson DG, Thibodeau SN, Waldman FM: Chromosomal instability in microsatellite-unstable and stable colon cancer. Clin Cancer Res 2006, 12: 6379–6385.

41. Chen W, Ding J, Jiang L, Liu Z, Zhou X, Shi D: DNA copy number profiling in microsatellite-stable and microsatellite-unstable hereditary non-polyposis colorectal cancers by targeted CNV array. Funct Integr Genomics 2017, 17: 85–96.

42. Ali H, Bitar MS, Al Madhoun A, Marafie M, Al-Mulla F: Functionally-focused algorithmic analysis of high resolution microarray-CGH genomic landscapes demonstrates comparable genomic copy number aberrations in MSI and MSS sporadic colorectal cancer. PLoS One 2017, 12: e0171690.

43. Sveen A, Johannessen B, Tengs T, Danielsen SA, Eilertsen IA, Lind GE, Berg KCG, Leithe E, Meza-Zepeda LA, Domingo E, et al: Multilevel genomics of colorectal cancers with microsatellite instability-clinical impact of JAK1 mutations and consensus molecular subtype 1. Genome Med 2017, 9: 46.

44. Cisyk AL, Nugent Z, Wightman RH, Singh H, McManus KJ: Characterizing Microsatellite Instability and Chromosome Instability in Interval Colorectal Cancers. Neoplasia 2018, 20: 943–950.

45. Roth A, Khattra J, Yap D, Wan A, Laks E, Biele J, Ha G, Aparicio S, Bouchard-Cote A, Shah SP: PyClone: statistical inference of clonal population structure in cancer. Nat Methods 2014, 11: 396–398.

46. Schell MJ, Yang M, Teer JK, Lo FY, Madan A, Coppola D, Monteiro AN, Nebozhyn MV, Yue B, Loboda A, et al: A multigene mutation classification of 468 colorectal cancers reveals a prognostic role for APC. Nat Commun 2016, 7: 11743.

47. Shin G, Greer SU, Xia LC, Lee H, Zhou J, Boles TC, Ji HP: Targeted short read sequencing and assembly of re-arrangements and candidate gene loci provide megabase diplotypes. Nucleic Acids Res 2019, 47: e115.

48. Bell JM, Lau BT, Greer SU, Wood-Bouwens C, Xia LC, Connolly ID, Gephart MH, Ji HP: Chromosome-scale mega-haplotypes enable digital karyotyping of cancer aneuploidy. Nucleic Acids Res 2017, 45: e162.

49. Greer SU, Nadauld LD, Lau BT, Chen J, Wood-Bouwens C, Ford JM, Kuo CJ, Ji HP: Linked read sequencing resolves complex genomic rearrangements in gastric cancer metastases. Genome Med 2017, 9: 57.

50. Xia LC, Bell JM, Wood-Bouwens C, Chen JJ, Zhang NR, Ji HP: Identification of large rearrangements in cancer genomes with barcode linked reads. Nucleic Acids Res 2018, 46: e19.

51. Li Y, Pang X, Cui Z, Zhou Y, Mao F, Lin Y, Zhang X, Shen S, Zhu P, Zhao T, et al: Genetic factors associated with cancer racial disparity - an integrative study across twenty-one cancer types. Mol Oncol 2020.

52. Chen X, Wang J, Mitchell E, Guo J, Wang L, Zhang Y, Hodge JC, Shen Y: Recurrent 8q13.2-13.3 microdeletions associated with branchio-oto-renal syndrome are mediated by human endogenous retroviral (HERV) sequence blocks. BMC Med Genet 2014, 15: 90.

53. Zare F, Dow M, Monteleone N, Hosny A, Nabavi S: An evaluation of copy number variation detection tools for cancer using whole exome sequencing data. BMC Bioinformatics 2017, 18: 286.

54. Lee SY, Chung H, Devaraj B, Iwaizumi M, Han HS, Hwang DY, Seong MK, Jung BH, Carethers JM: Microsatellite alterations at selected tetranucleotide repeats are associated with morphologies of colorectal neoplasias. Gastroenterology 2010, 139: 1519–1525.

55. Mandal R, Samstein RM, Lee KW, Havel JJ, Wang H, Krishna C, Sabio EY, Makarov V, Kuo F, Blecua P, et al: Genetic diversity of tumors with mismatch repair deficiency influences anti-PD-1 immunotherapy response. Science 2019, 364: 485–491.

56. Plaschke J, Kruger S, Jeske B, Theissig F, Kreuz FR, Pistorius S, Saeger HD, Iaccarino I, Marra G, Schackert HK: Loss of MSH3 protein expression is frequent in MLH1-deficient colorectal cancer and is associated with disease progression. Cancer Res 2004, 64: 864–870.

57. Forgo E, Gomez AJ, Steiner D, Zehnder J, Longacre TA: Morphological, immunophenotypical and molecular features of hypermutation in colorectal carcinomas with mutations in DNA polymerase epsilon (POLE). Histopathology 2020, 76: 366–374.

