## supplementary_materials for "Profiling diverse sequence tandem repeats in colorectal cancer reveals co-occurrence of microsatellite and chromosomal instability involving Chromosome 8"

**TITLE**

### SUPPLEMENTARY METHOD

#### Primer design

Using a single base offset, we generated all of the 20-mer sequences from the GRCh37.1 human genome reference. We determined the alignment of these 20-mers in terms of their unique or repetitive location in the human genome and accounted for exact matches with a tolerance for either one or two mismatches. To identify targeting primers, we used several criteria: (i) primer probes upstream and downstream of any given target for double stranded coverage of the targeted region within 100 to 200 bases; (ii) adjustable primer probe density; (iii) a primer probe GC content between 30% and 65%; (iv) uniqueness of the last 20 bases for the 3' portion of the primer probe in the human genome with a string edit distance of 1 from any other genome location; (v) no overlap of the last 10 bases with a known SNP as annotated in dbSNP Build ID 131; and (vi) no sequences immediately adjacent to highly repetitive sequences.

#### Sequencing of microsatellite and genomic targets

The top (5'- CGAGATCTACACTCTTTCCCTACACGACGCTCTTCCGATCxxxxxx*T), which contains a phosphorothioate bond (indicated by *), and bottom (5'-/5Phos/ xxxxxxGATCGGAAGAGCGTCGTGTAGGGAAAGAGTGTAGATCTCG) multiplex adapters are standard desalted ultramer oligonucleotides (Integrated DNA Technologies, Coralville IA). These adapters contain a 7-base indexing sequence (xxxxxx*T) directly following the sequencing primer binding site. The adapters were annealed in a final concentration of 15 µM per adapter in Nuclease Free Duplex Buffer (IDT) by a 1% temperature ramp from 94°C to 20°C, after an initial 5 min 94°C denaturation step. Unlike standard Illumina adapters, our modified library adapters are only complementary to the P5 primer on the flow cell surface. The portion that is complementary to the P7 primer is introduced in the primer probe extension step. Primer probes were column-synthesized at the Stanford Genome Technology Center and combined in an isomolar pool.

The KAPA HyperPlus library preparation kit (Roche) was used for the following steps. For each library, 1 μg gDNA was subject to random fragmentation with the KAPA enzyme mix; the incubation was at 37°C for 9 min, directly followed by incubation on ice. A-tailing enzyme mix was added to the final fragmentation products and the fragmented library was A-tailed with incubation at 65°C for 30 min. Because the random fragmentation creates blunt-ended breaks, the end-repair step was omitted. The DNA ligase mix including 75 pmol annealed adapter was added to the A-tailed library. The reaction volume was incubated at 20°C for 15 min. Afterwards, the library products were purified with AMPure XP beads in a bead solution-to-sample ratio of 0.8. The size distribution of the sequencing library was measured with the DNA High Sensitivity Reagent Kit on a LabChip GX (Perkin-Elmer) per the manufacturer's protocol. The purified library was then used directly for single primer targeted sequencing with no additional steps.

The flow cell modification and capture assay procedures were modified from those previously described [23]. The targeting process requires (1) hybridization and extension of the target oligonucleotides onto a solid phase support such as a flow cell, followed by capturing of the sequencing library by overnight hybridization; and (2) extension of the captured library and standard Illumina cluster generation. For runs performed using High Output mode, we used v3 sequencing reagents (Illumina) for 2x101 cycle paired end reads. For runs performed using Rapid Run mode, we used v2 reagents (Illumina) for 125 (read 1) and 101 (read 2) cycles of paired end sequencings. For all the HiSeq experiments, image analysis and base calling were performed using the HCS 2.2.58 and RTA 1.18.64 software (Illumina).

**Digital PCR confirmation of copy number alterations**

For a number of amplifications, we used digital PCR assay for validation. First, we identified the coordinates defining individual gene regions of interest (ROI) within their exons. We designed all copy number primers and probes with Primer3, using input gene sequences from NCBI Nucleotide's GRCh37.p10 Primary Assembly [27, 28]. Each primer was verified for uniqueness with UCSC Genome Browser's Blat based on the GRCh37/hg19 assembly [29]. We obtained all primers from Integrated DNA Technologies (San Diego, CA) and MGB hydrolysis probes from Life Technologies (NY, USA). For the genes *ERBB2*, *FGFR1* and *MET,* we used a hydrolysis probe-based primer design, including a FAM-tagged MGB probe flanked by two primers for each ROI. We used primers and a VIC-probe for a previously defined [20] highly conserved region on Chromosome 1 (UC1) as a reference for all copy number assays. This method has been previously validated for detecting amplifications in cancer [19, 20].

For the genes *AURKA*, *CDK4*, *FLT3* and *VEGFA,* we utilized an analogous multiplex digital PCR assay using a single-color DNA-binding dye (EvaGreen) as we have previously described [30]. Briefly, the multiplexing is based on discriminating differences in fluorescence intensity as dependent on amplicon length. In our previous work, we showed the ability to discriminate amplicons that differed by as little as 10bp. Here we use the same principle to multiplex an ROI and a reference, with ROI primers amplifying similar or slightly longer amplicons (60-66 bp). The droplets can then be clustered based on the difference in fluorescent signal amplitude; populations containing no template correspond to the lowest fluorescent amplitude. This design allowed us to multiplex the ROI and UC1 assays in the same well.

The digital PCR assays were run on a QX200 droplet digital PCR system (Bio-Rad) across a series of annealing temperatures [19]. The 20 µl single-color ddPCR reaction mixtures consisted of 2X ddPCR Evagreen Supermix, 100 nM each primer and 20 ng restriction digested gDNA. Droplets were generated as described above and cycled as follows: 95ºC for 5 min (1 cycle); 95ºC for 30s, 52-62ºC for 1 min (40 cycles); 4ºC for 5 min, 90 ºC for 5 min (1 cycle); and 4ºC hold. After PCR, each well was read individually with the QX200™ droplet reader on one fluorescent channel. We identified the annealing temperature that produced four distinct and fully separated droplet populations: droplets containing no template, reference amplicon only droplets, target amplicon only droplets, or both the reference and target amplicon template fragments.

We assessed each patient sample with three independent replicates for gene copy number. For samples analyzed in the hydrolysis probe-based assay, droplets were clustered using QuantaSoft (version1.2.10.0). For samples analyzed with the single-color dye-based assay, we exported the raw droplet fluorescence amplitudes from Bio-Rad QuantaSoft software. The concentration of each target was calculated as follows:

$$\frac{-\ln(\frac{negative\ droplets}{total\ droplets})}{droplet\ volume}$$

where *droplet volume* is the volume of an individual droplet in µls. We set this number as 0.0009ul for EvaGreen assays. Copy number of each well was calculated as follows:

$$2 * \left(\frac{[ROI]}{[UC1]}\right)$$

We calculated a weighted average and standard deviation of copy number across triplicates wells, based on the total number of droplets read in each replicate. We called a sample with a copy number of 3-5 copies/genome a gain and a sample with > 5 copies a high-level amplification.  We defined loss as copy number ≤ 1, corresponding to a loss of one to two copies.

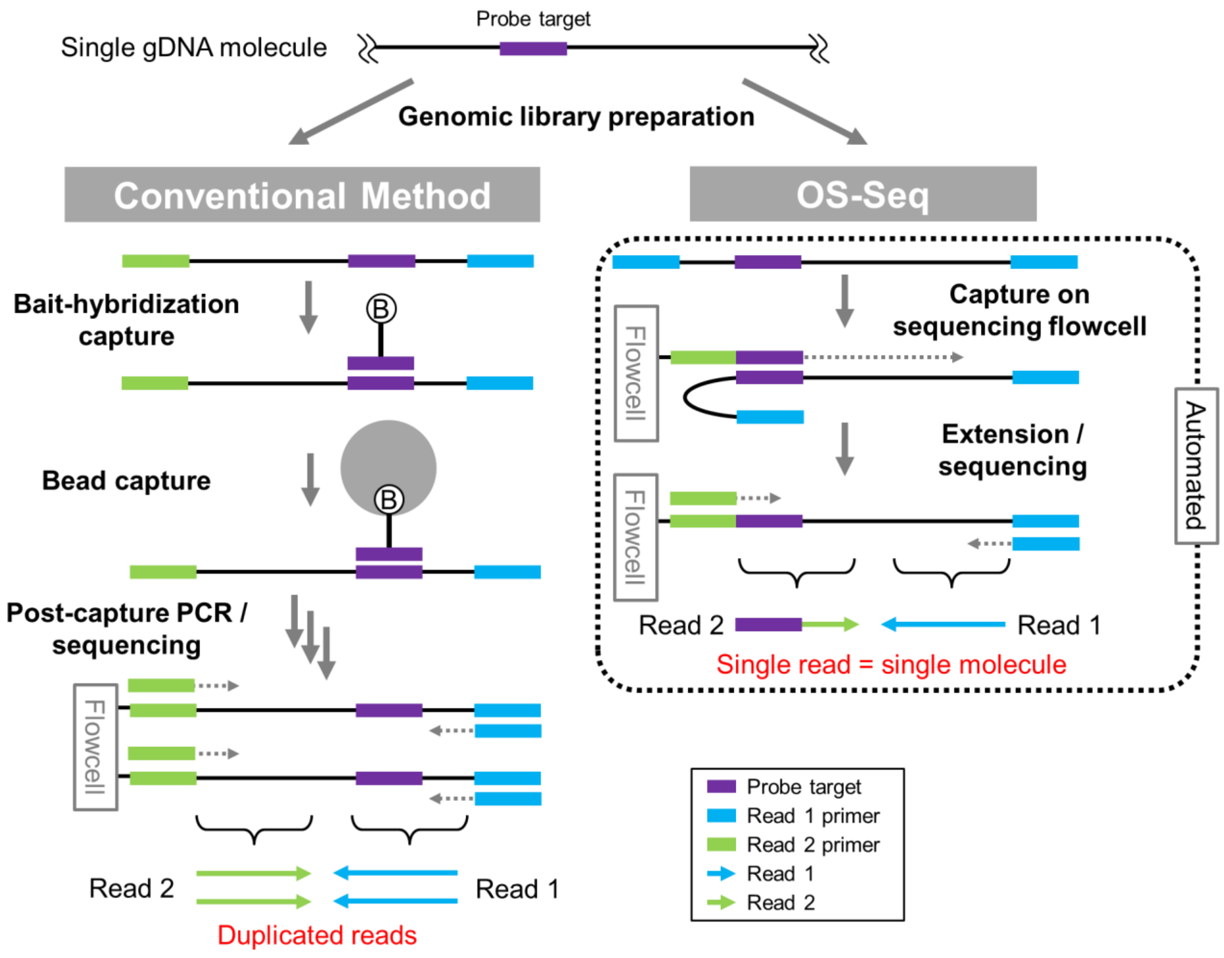


**Supplementary Figure S1. Comparison between conventional and single primer targeted sequencing methods.**
The single primer-targeted sequencing method used in the current study (OS-Seq) is compared with a conventional method, generally used for the whole exome and gene panels. This single primer targeting method uses a single adapter library, while the conventional method uses a ready-to-sequence dual-adapter library. The capture process itself completes the libraries for the subsequent sequencing step. Steps for targeting genomic sequence involves a round of hybridization and single primer extension. This process is simpler compared to the conventional method which generally uses multiple wash steps after immobilizing the captured library with magnetic beads. Moreover, targeted sequencing is easily automated with a commercial system (e.g. Illumina cBot). Most importantly, OS-Seq requires no post capture PCR step when using capture probes already immobilized on the sequencing flowcell. Therefore, a sequencing read represents a single DNA molecule. PCR-born indels, or stutters can be minimized especially when targeting microsatellites.

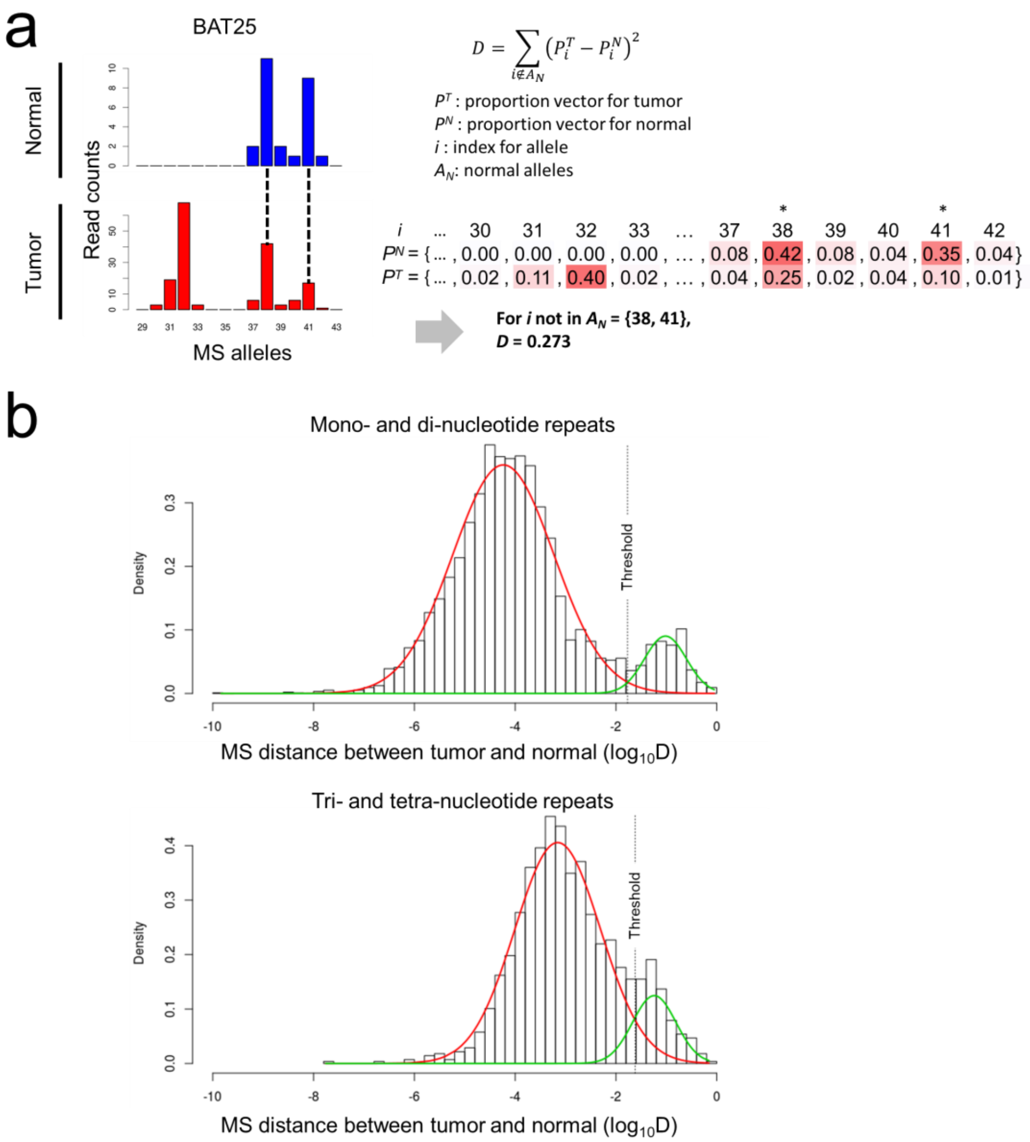


| $i$ | ... | 30 | 31 | 32 | 33 | ... | 37 | 38* | 39 | 40 | 41* | 42 |
|---|---|---|---|---|---|---|---|---|---|---|---|---|
| $P^N$ = { | ... | 0.00 | 0.00 | 0.00 | 0.00 | ... | 0.08 | 0.42 | 0.08 | 0.04 | 0.35 | 0.04} |
| $P^T$ = { | ... | 0.02 | 0.11 | 0.40 | 0.02 | ... | 0.04 | 0.25 | 0.02 | 0.04 | 0.10 | 0.01} |

**Supplementary Figure S2. Determination of microsatellite allelic shift.**
**(a)** Definition of microsatellite distance between tumor and normal samples. An example of allelic shift at the BAT25 locus of the P799 tumor is shown. Left panels show the allele histograms generated by our OS-Seq method. Relative abundance (sequencing read count) of DNA molecules and different microsatellite alleles (number of motif repeats) are indicated on the y- and x-axes, respectively. From the normal allele profile, the heterozygote alleles (38 and 41 motif repeats) are apparent. The allele proportion is calculated by dividing the read count of each allele by the total read count from the locus. The example shown at the bottom right are the proportion vectors from the normal and tumor samples ($P^N$ and $P^T$), and the distance value calculated following the definition of microsatellite distance provided at the top right. **(b)** Kernel density distribution of microsatellite distance values from all 46 tumor / normal comparisons at 225 microsatellite loci. The distribution is separately shown for mono- / dinucleotide and tri- / tetranucleotide microsatellites. Mixture of two Gaussian distributions are assumed: one for wild type, and the other for shifted alleles. A threshold (dotted vertical line) is indicated at the intersection of the two density curves.

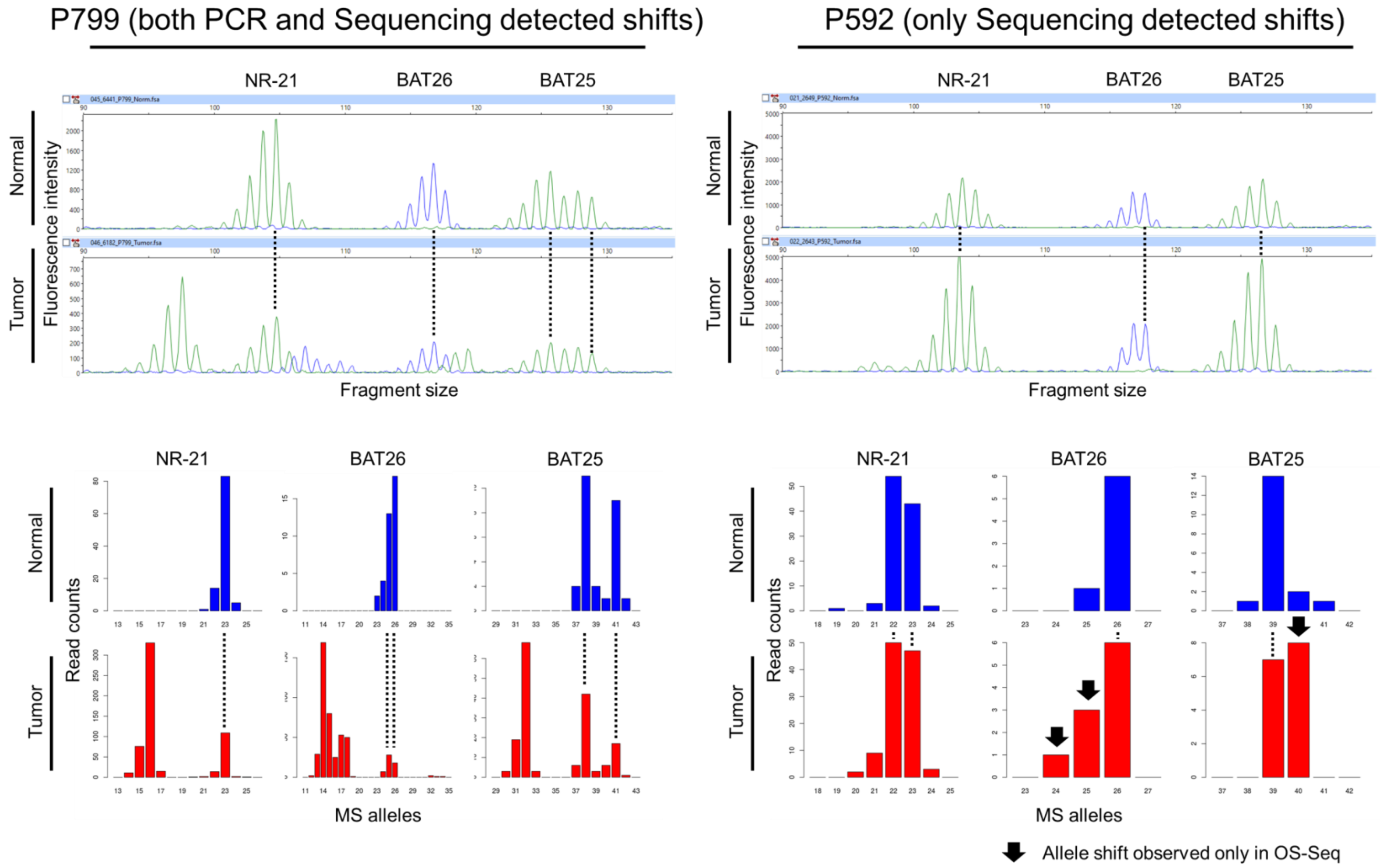


**Supplementary Figure S3. Microsatellite allelic shifts detected by PCR-CE and OS-Seq methods.**
For three microsatellite loci (NR-21, BAT26, and BAT25), allele profiles of both normal and tumor samples are shown for two examples (P799 and P592 cases). The P799 and P592 tumors are classified as MSI and MSS according to our sequencing analysis. Electropherograms generated by PCR-CE method (top panels) provide relative abundance (y-axis, fluorescence intensity) of amplicons with different sizes (x-axis, DNA size in bp). Although it has erroneous stutter amplification, the tumor allele profile from the P799 tumor (MSI) suggests allele shifts. On the other hand, the tumor allele profile from the P592 tumor (MSS) shows no change compared to the normal profile. The allele histograms generated by our targeted sequencing analysis method (bottom panels) provide relative abundance (y-axis, sequencing read count) of DNA molecules, including different microsatellite alleles (x-axis, number of motif repeats). The microsatellite profiles show dramatically decrease stutter amplification compared to the profiles generated by PCR-CE method. For the P799 tumor, the PCR-CE and targeted sequencing profile match. However, for the P592 tumor, the sequencing profile show allele shits at BAT26 and BAT25 loci (black arrows), which are invisible from the PCR-CE profile due to the stutter errors.

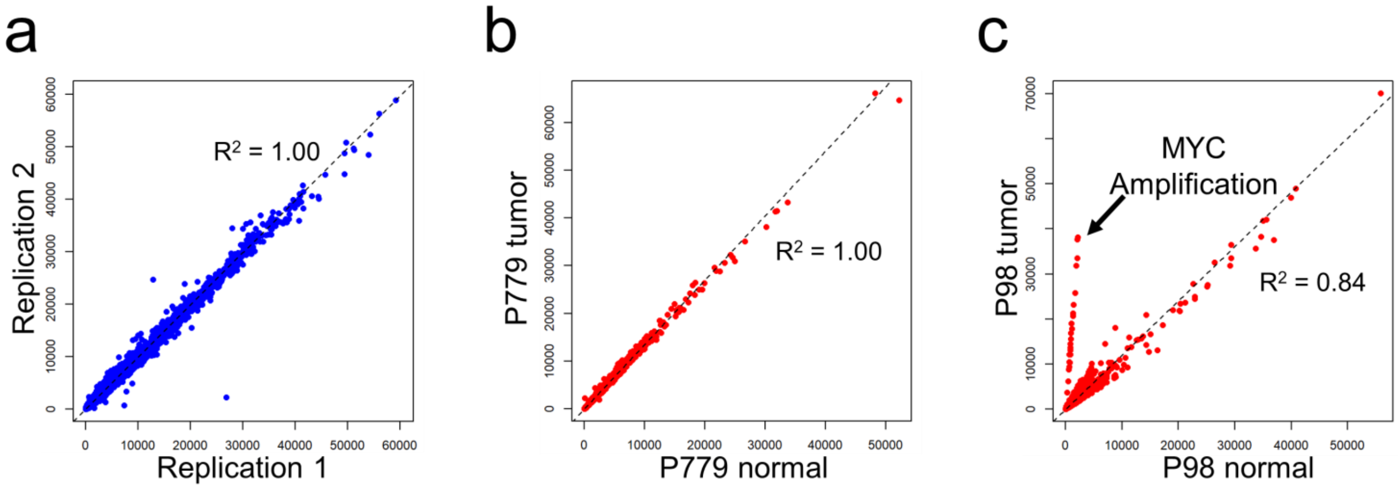


**Supplementary Figure S4. Detection of copy number aberration by using reproducible target capture.**
**(a)** Reproducibility of capture efficiency between replications. All tumor samples were sequenced by two independent library preparations labeled by different sample barcodes. Every sequencing read is tagged by an index corresponding to the capture primer-probe. Read counts per each index are compared in the two replications for all 46 tumor samples. **(b)** Reproducibility of capture efficiency between different samples. Read counts per each index are compared in normal and tumor samples from P779, among which the tumor sample does not have copy number instability (e.g. CS tumors). **(c)** An example of focal amplification. Read counts per each index are compared in normal and tumor samples from P98. A group of indexes have shared a ratio 12-fold greater than the average, and all of them target *MYC*. In all plots, black dashed line indicates linear regression, and the correlation is indicated as R-squared value.

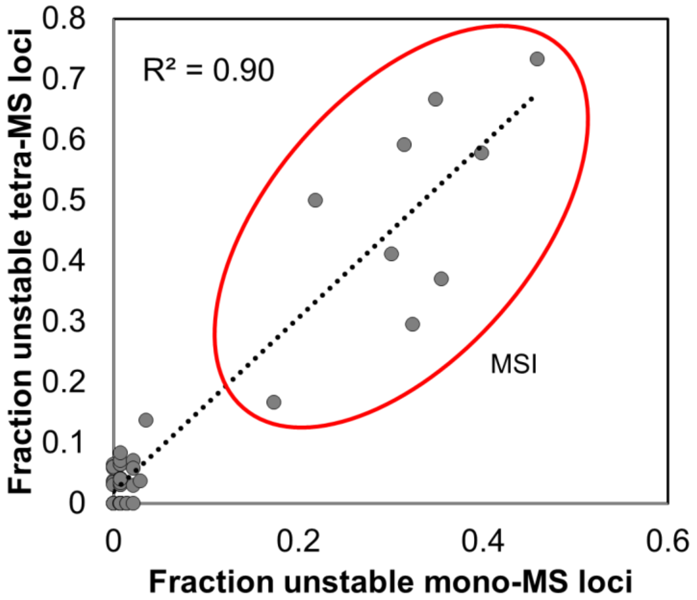


**Supplementary Figure S5. Correlation between fractions of unstable mono- and tetranucleotide microsatellite loci.**

For the two microsatellite classes (mono- and tetranucleotide repeats), the scatter plot compares fractions of unstable microsatellite loci.  The fraction of instability in each class was calculated by dividing the number of somatic microsatellite allelic shifts by the total number of genotyped microsatellites.  Black dotted line indicates linear regression, and the correlation is indicated as R-squared value.  MSI tumors are clustered (red oval) as they have an elevated instability in both the microsatellite classes.

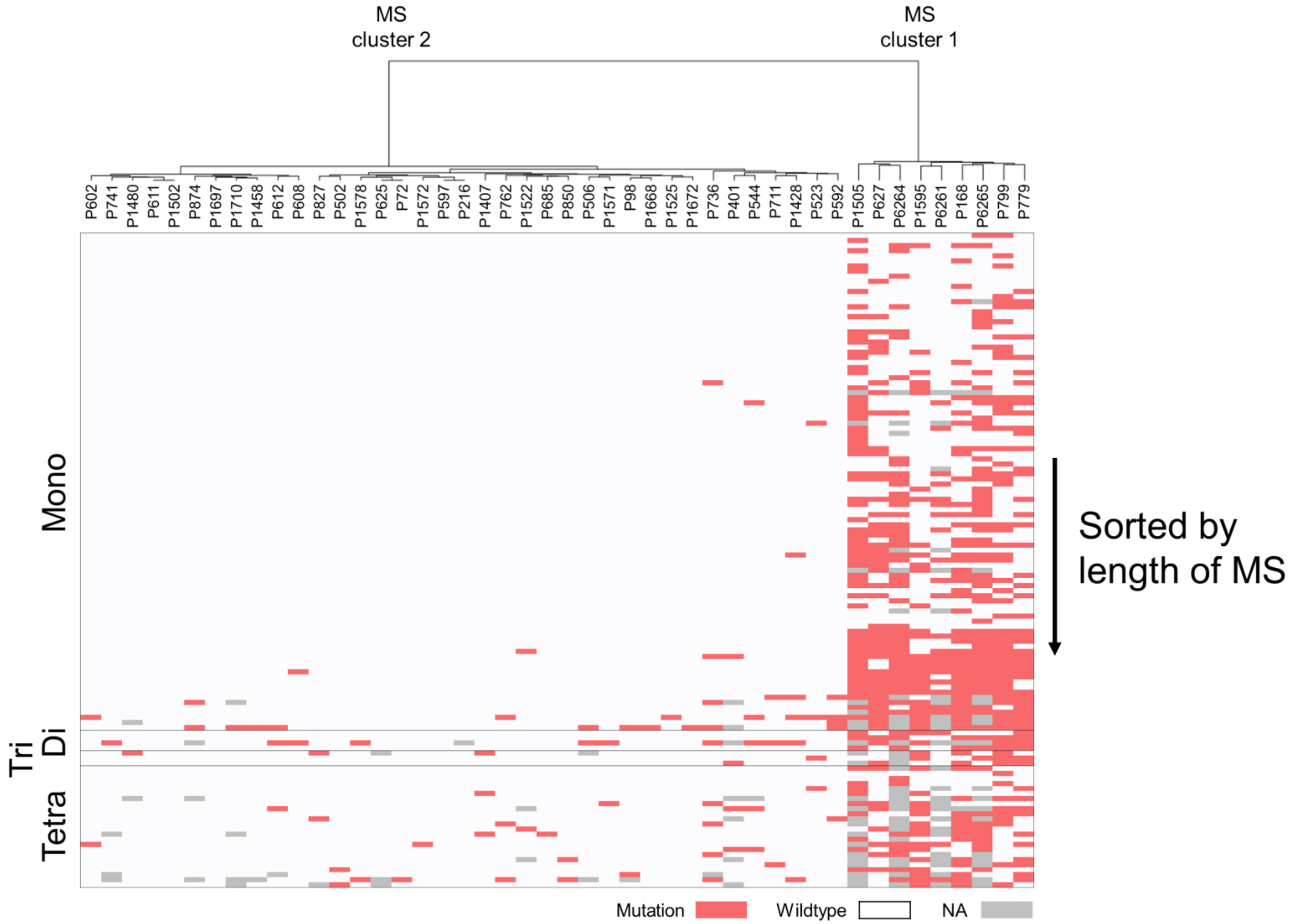


**Supplementary Figure S6. Hierarchical clustering based on mutation profile of 225 microsatellites.**
Clustering based on 225 microsatellites across four different classes. A 225x46 matrix including the presence (1) or absence (0) of microsatellite allele shift mutation was used for an unsupervised hierarchical clustering, which generated two clusters (MS Clusters 1 and 2). The heatmap shown here is matrix for a subset (129 microsatellites), which had at least one allelic shift mutation among the 46 tumors. 'NA' indicates undetermined due to no or limited sequencing coverage.

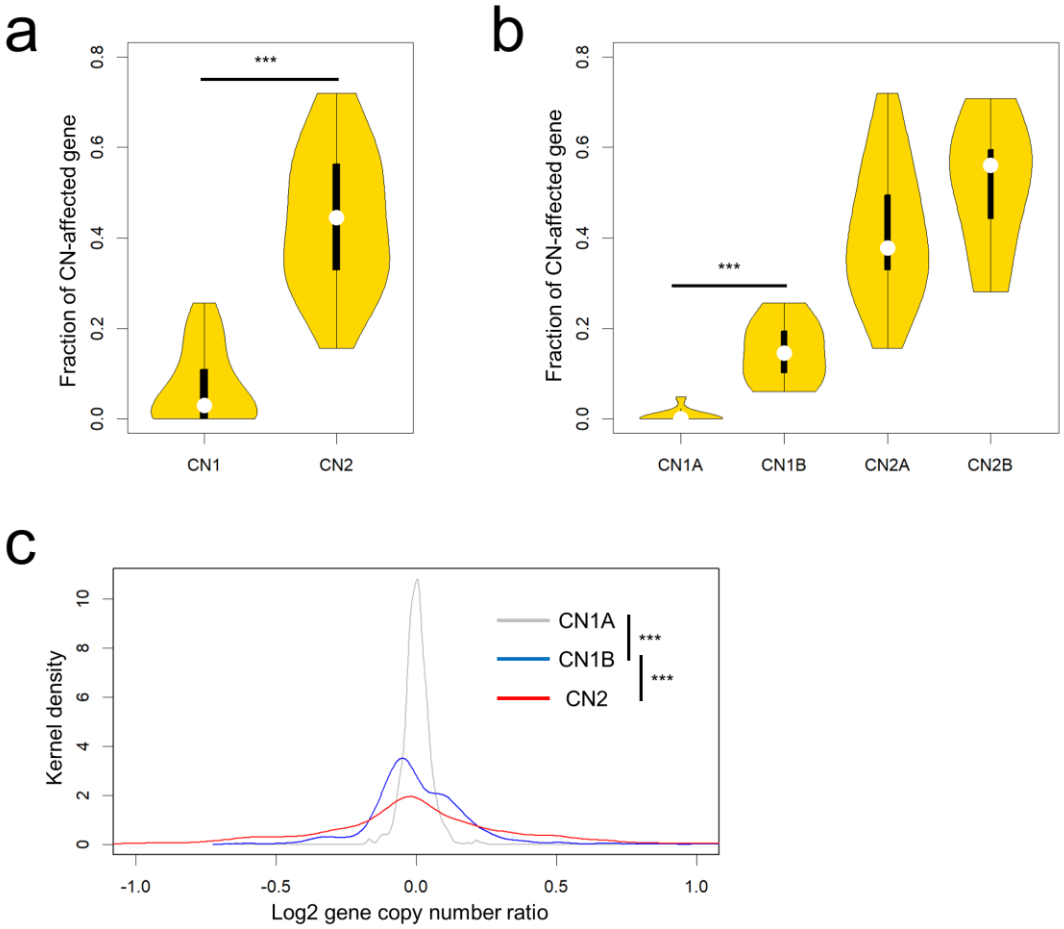


**Supplementary Figure S7. Validation of hierarchical clustering based on copy number profile.**
**(a and b)** Comparison of fraction of affected genes among different CN clusters. The fraction of copy number-affected genes was significantly different ($p < 0.001$) between the two major clusters (CN-1 and CN-2), and between CN-1A and CN-1B clusters. The violin plot shows normalized density distribution of among the tumor samples, and the white dot indicates the median value. **(c)** Comparison of variance of gene copy number ratio. The Kernel density distributions for CN-1A, CN-1B, and CN-2 clusters are shown, of which the variances are significantly different ($p < 0.001$ per F-test).

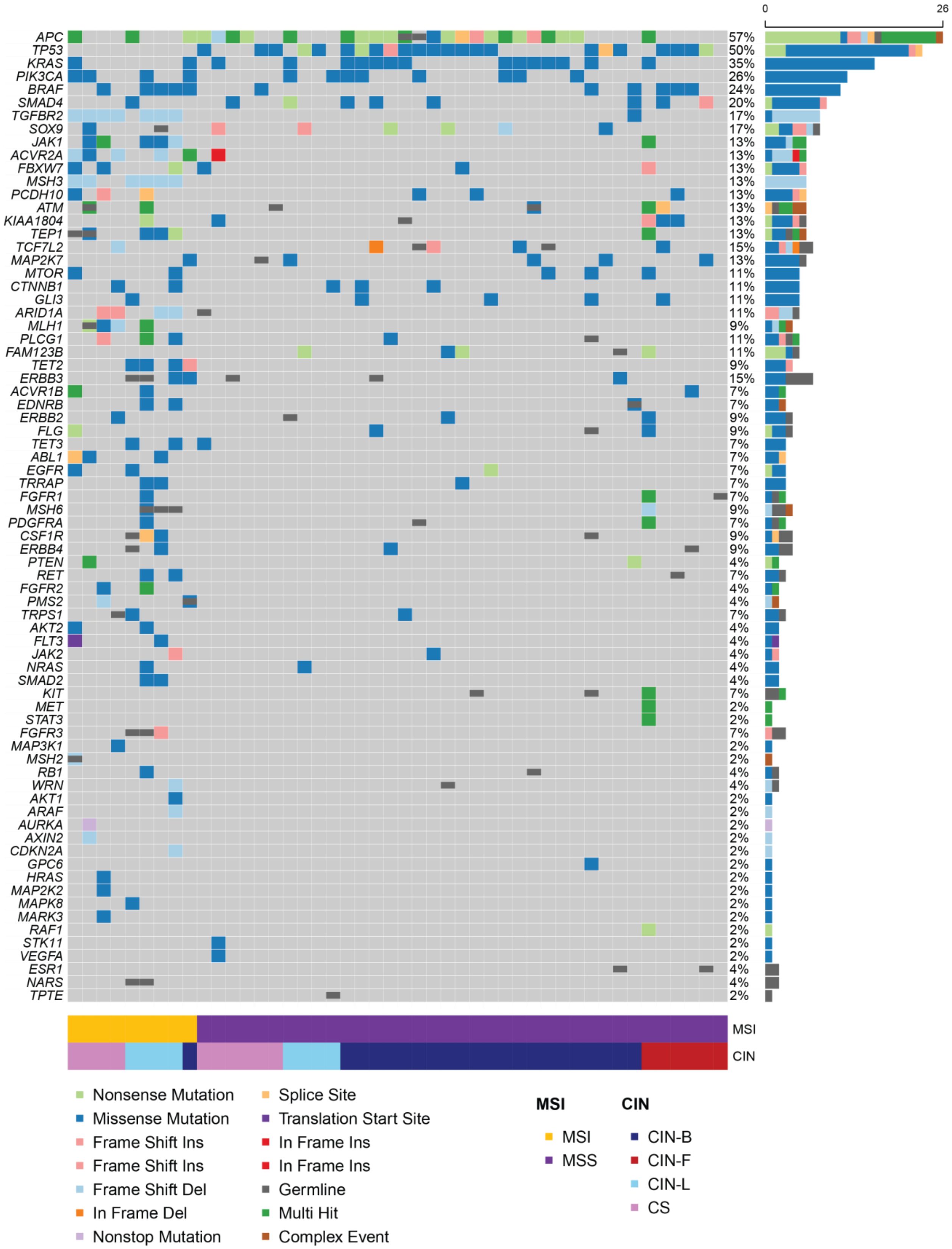


**Supplementary Figure S8. Oncoplot for all the target driver genes.**
For all the target genes (rows), mutation profiles of 46 tumors (columns) are shown. Only mutations with a CADD score greater than 20 are used. Different types of somatic mutations are shown as squares with different colors. Gray color indicates no mutation call at the given gene. Germline mutations are also indicated with a smaller rectangle overlaid on the somatic mutation map. Right panel shows the number of affected samples for each gene. Genes are sorted according to the frequency of somatic mutations. Lower panel indicates MSI and CIN sample annotations determined by the sequencing assay.

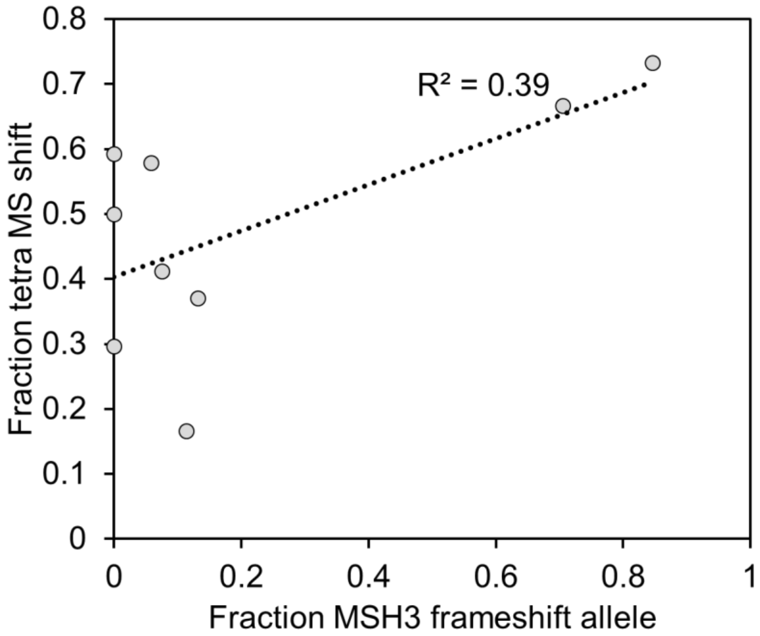


**Supplementary Figure S9. Association between a *MSH3* frameshift and EMAST.**
For the nine MSI samples, fraction of MSH3 frameshift allele and degree of EMAST (i.e. fraction of unstable tetranucleotide microsatellite loci) are compared. Black dotted line indicates linear regression, and the correlation is indicated as R-squared value.

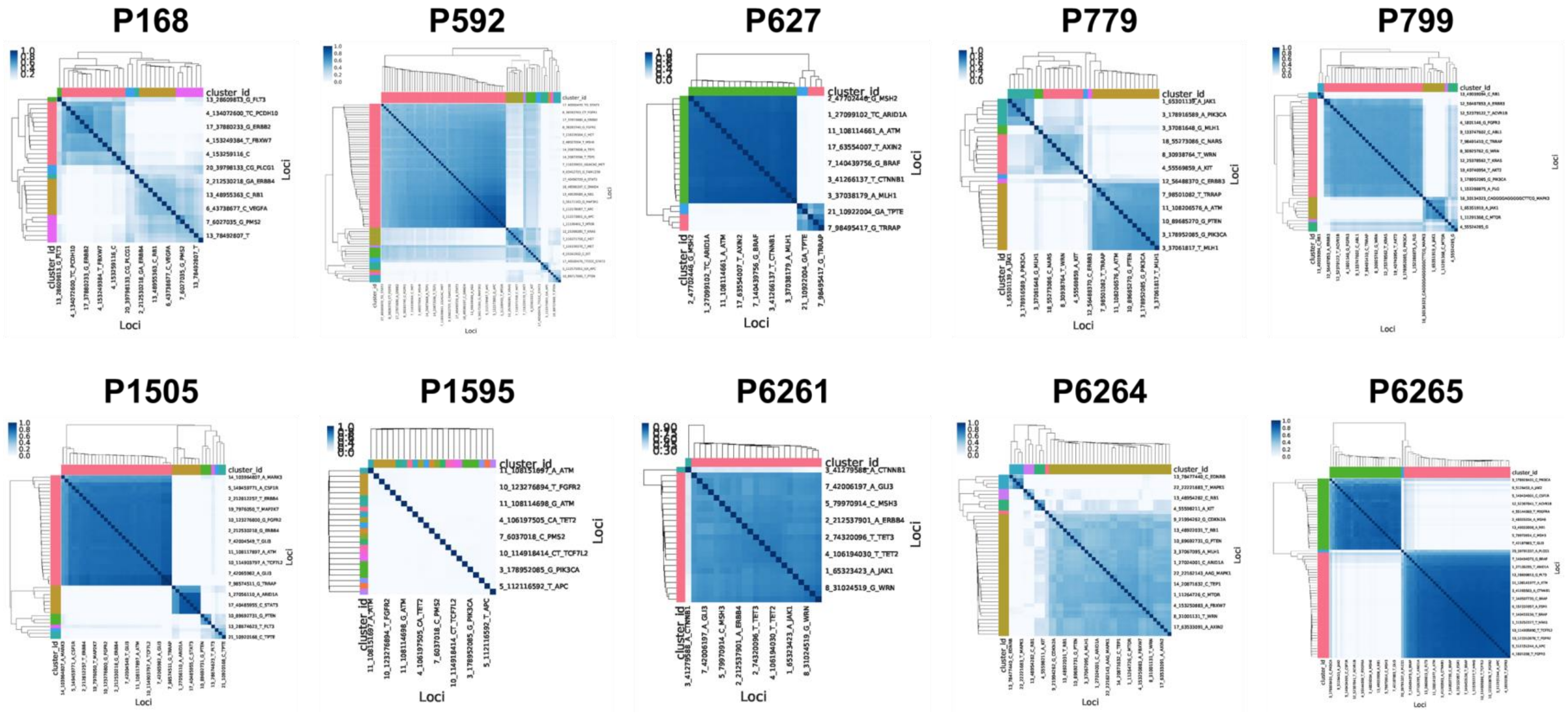


**Supplementary Figure S10. PyClone clustering of somatic mutations.**
Similarity matrixes are shown with PyClone clustering results. The heat map indicate distance of estimated cellular prevalence between two mutations. The mutations in clusters are indicated by different colors at the first row and the first column of each matrix.

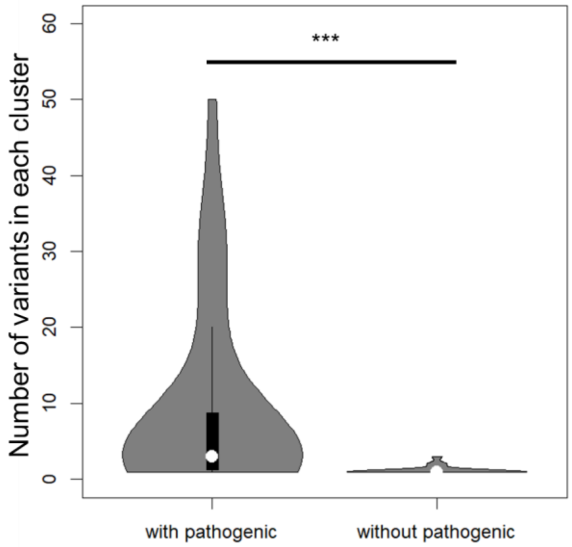


**Supplementary Figure S11. Distribution of PyClone cluster size.**
Distribution of the Pyclone cluster size (the number of mutations in each cluster) is significantly different ($p < 0.001$) between the clusters with and without a pathogenic mutation (>CADD-20). Violin plot shows normalized density distribution among the clusters, and white dot indicates the median value.

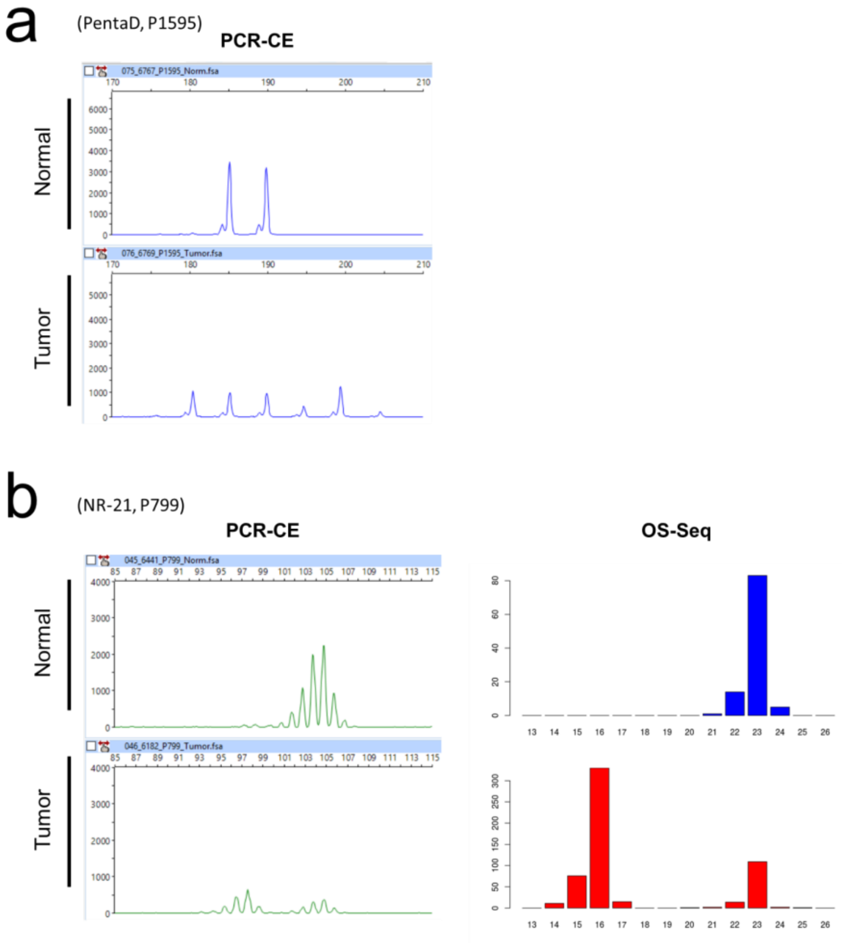


**Supplementary Figure S12. Microsatellite analysis results supporting subclonal structure of MSI tumors**

**(a)** Six tumor alleles at a microsatellite loci. For a pentanucleotide microsatellite locus (PentaD), allele profiles of both normal and tumor samples are shown for P1595. Electropherograms generated by PCR-CE method provide relative abundance (y-axis, fluorescence intensity) of amplicons with different sizes (x-axis, DNA size in bp). The tumor allele profile shows six different microsatellite alleles including the two matching normal alleles. **(b)** An example of relatively homogeneous tumor microsatellite alleles. For a microsatellite locus (NR-21), allele profiles of both normal and tumor samples are shown for P799. Electropherograms generated by PCR-CE method are shown in left panels. The allele histograms generated by our targeted microsatellite profiling (right panels) also provide relative abundance (y-axis, sequencing read count) of DNA molecules, including different microsatellite alleles (x-axis, number of motif repeats). Both methods are supportive to each other, although our sequencing approach has a reduced degree of stutter artifacts.

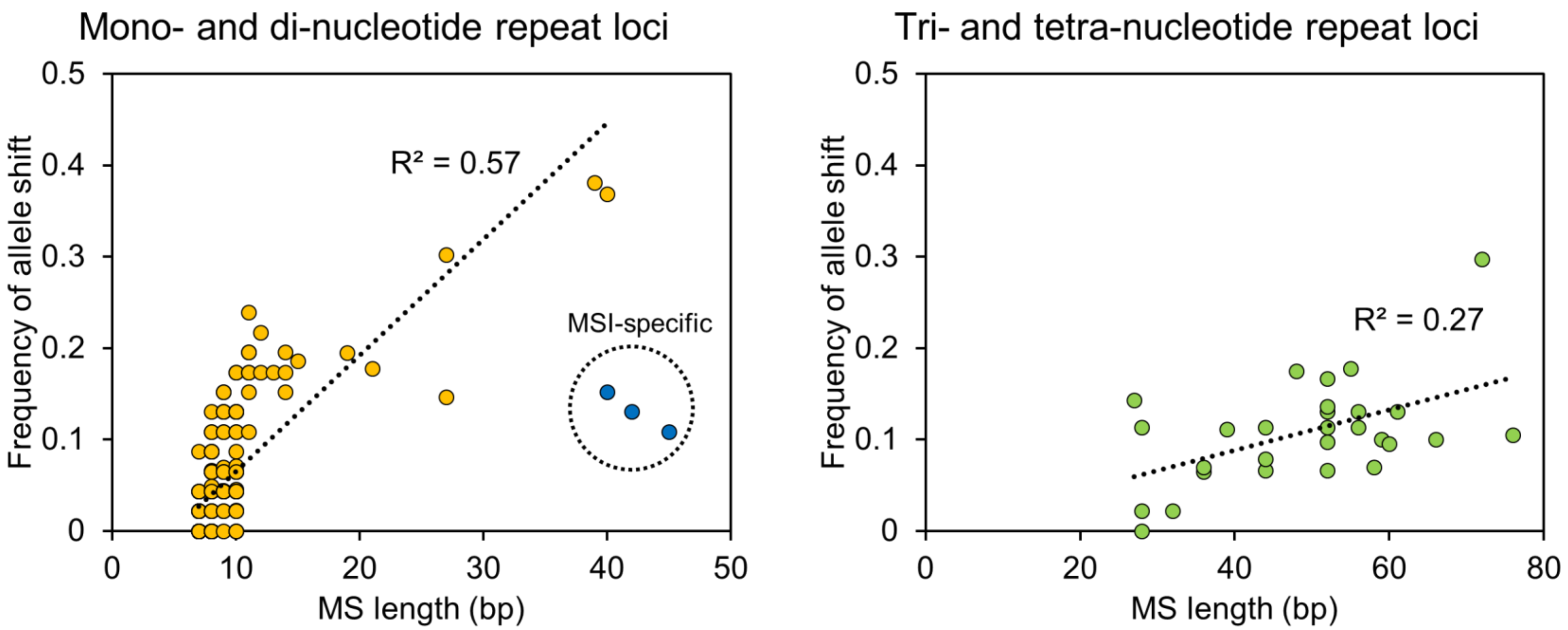


**Supplementary Figure S13. Correlation between length of microsatellite and mutation frequency.**
The four microsatellite classes are grouped into two by their motif length (i.e. shorter and longer). For both the groups, the length of microsatellite are positively correlated with the frequency of mutation among the 46 CRCs. Black dotted line indicates linear regression, and the correlation is indicated as R-squared value. For mono- and dinucleotide microsatellites, there were three outliers (dotted circle) with relatively low mutation frequency compared to other microsatellites with similar length. These were specific to MSI tumors. Two of them were dinucleotide microsatellites in the Bethesda panel (D2S123 and D5S346), and the other was a long homopolymer but with many interruptions.
